# XAB2 prevents abortive recombinational repair of replication-associated DNA double-strand breaks and its loss is synthetic lethal with RAD52 inhibition

**DOI:** 10.1101/2020.04.10.035410

**Authors:** Abhishek B. Sharma, Hélène Erasimus, Lia Pinto, Marie-Christine Caron, Katrin Neumann, Petr V. Nazarov, Barbara Klink, Sabrina Fritah, Christel C. Herold-Mende, Simone P. Niclou, Patrick Calsou, Jean-Yves Masson, Sébastien Britton, Eric Van Dyck

**Affiliations:** DNA Repair and Chemoresistance Group, Department of Oncology, Luxembourg Institute of Health (LIH), Luxembourg, Luxembourg; Faculty of Science, Technology and Communication, University of Luxembourg, Esch-sur-Alzette, Luxembourg; CHU de Québec Research Center, Oncology Division, Québec City, Canada; Department of Molecular Biology, Medical Biochemistry and Pathology, Laval University Cancer Research Center, Québec City, Canada; Quantitative Biology Unit, Multiomics Data Science Group, LIH, Luxembourg, Luxembourg; National Center of Genetics, Laboratoire National de Santé, Dudelange, Luxembourg; Functional Tumour Genetics Group, Department of Oncology, LIH, Luxembourg; NorLux Neuro-Oncology Laboratory, Department of Oncology, LIH, Luxembourg; Department of Neurosurgery, University Clinic Heidelberg, Heidelberg, Germany; Department of Biomedicine, University of Bergen, Norway; Institut de Pharmacologie et de Biologie Structurale, Université de Toulouse, CNRS, UPS, Toulouse, France, Equipe Labellisée Ligue Nationale Contre le Cancer 2018

**Author notes:** These authors contributed equally (Abhishek B. Sharma, Hélène Erasimus). These authors contributed equally (Patrick Calsou, Jean-Yves Masson, Sébastien Britton).

**Keywords:** DNA double-strand break (DSB) repair, single-ended DNA double-strand break (seDSB), collapsed replication fork, homologous recombination, non-homologous end-joining, DNA end resection, XAB2, RAD52, RAD51, Ku, synthetic lethality, glioblastoma, temozolomide, camptothecin, chemotherapy

## Abstract

Unrepaired O^6^-methylguanine lesions induced by the alkylating chemotherapy agent temozolomide lead to replication-associated single-ended DNA double-strand breaks (seDSBs) that are repaired predominantly through RAD51-mediated homologous recombination (HR). Here, we show that loss of the pre-mRNA splicing and DNA repair protein XAB2 leads to increased temozolomide sensitivity in glioblastoma cells, which reflects abortive HR due to Ku retention on resected seDSBs. XAB2-dependent Ku eviction also occurred at seDSBs generated by the topoisomerase I poison campthotecin and operated in parallel to an ATM-dependent pathway previously described. Although Ku retention elicited by loss of XAB2 did not prevent RAD51 focus formation, the resulting RAD51-ssDNA associations were unproductive, leading to increased engagement of non-homologous-end-joining in S/G2 and genetic instability. Overexpression of RAD51 or the single-stranded DNA annealing factor RAD52 rescued the XAB2 defects. RAD52 depletion led to severe temozolomide sensitivity, whereas a synthetic lethality interaction was observed between RAD52 and XAB2.

## Introduction

Despite surgical resection, ionizing radiation and chemotherapy with the DNA alkylating agent temozolomide (TMZ), glioblastoma (GBM) remains one of the most aggressive and lethal cancers [1, 2], due to chemoresistance driven by complex DNA repair mechanisms. The DNA repair protein O^6^-methylguanine-DNA methyltransferase (MGMT) removes the most cytotoxic lesion induced by TMZ, O^6^-methylguanine (O^6^-meG) [3], providing a major TMZ resistance mechanism [4]. Epigenetic silencing of *MGMT* is a predictive and prognostic biomarker in TMZ-treated GBM patients [5]. O^6^-meG lesions left unrepaired by MGMT generate O^6^-meG/thymidine mismatches during S phase. These mismatches are recognized, but not resolved, by the mismatch repair (MMR) pathway, resulting in futile repair cycles and persistent single-stranded DNA (ssDNA) gaps that cause replication fork collapse and DNA double-stranded DNA breaks (DSBs) during a subsequent round of replication [6, 7].

DSBs resulting from replication fork collapse are single-ended DSBs (seDSBs) [8]. As for two-ended DSBs, seDSBs can be processed by homology-directed or end-joining mechanisms. Homologous recombination (HR) mediated by the RAD51 recombinase plays a central role in replication fork repair in mammalian cells [9] through a pathway called break-induced replication (BIR) [10]. HR is involved in the repair of lesions resulting from O^6^-meG adducts [11, 12]. During BIR, the seDSB is resected to provide Replication Protein A (RPA)-coated, 3’ ssDNA overhangs on which RAD51 operates to replace RPA and assemble nucleoprotein filaments. These filaments mediate homology search and strand invasion into the homologous sister chromatid, generating a displacement loop (D-loop) [13]. The single-strand annealing (SSA) factor RAD52 facilitates the assembly of ssDNA/RAD51 nucleoprotein filaments during HR-mediated seDSB repair in human cells [14]. Similar to yeast RAD52 [13], human RAD52 can also promote BIR mechanisms without RAD51. Thus, in cancer cells undergoing replication stress, a BIR pathway has been described which critically depends on RAD52 [15]. RAD52-mediated BIR also promotes mitosis DNA synthesis (MiDAS) at common fragile sites, a process where RAD51 is dispensable [16]. RAD52 forms ring structures that interact with ssDNA, duplex DNA, and DSB ends [17-19]. RAD52 rings expose ssDNA at their outer surface [18, 20, 21] and catalyse the annealing of complementary strands generated during DSB end resection [22-24], as well as second-end capture in the repair of double-ended DSBs [25, 26]. Relevant to BIR, human RAD52 also promotes DNA strand exchange and D-loop formation in vitro [27-29].

Processing of seDSBs by non-homologous end-joining (NHEJ) is a toxic mechanism as it involves the juxtaposition and ligation of distant DNA ends, resulting in chromosomal aberrations and genetic instability [30]. Unlike HR, which is restricted to the S and G2 phases of the cell cycle, NHEJ is active throughout interphase, including G1. In addition, seDSBs termini are initially sequestered by the DNA end-binding heterodimer Ku, a crucial NHEJ factor [31]. Ku binding at seDSBs promotes NHEJ [32] and impairs RAD51-mediated HR [33]. However, fully active HR outcompetes NHEJ in repairing seDSBs in S phase [32].

DNA end resection occurs in the S and G2 phases of the cell cycle and involves the MRE11-RAD50-NBS1 (MRN) complex, C-terminal-binding protein interacting protein (CtIP), exonuclease 1 (EXO1), Bloom syndrome protein (BLM), and DNA2 nuclease/helicase [34]. Resection is initiated at some distance from the seDSB by a nick introduced by the endonuclease activity of MRE11, itself activated by ataxia telangiectasia-mutated (ATM) kinase and CtIP. Bidirectional resection then takes place, mediated by MRE11 exonuclease activity in the 3’-5’ direction and EXO1/BLM/DNA2 in the opposite direction, generating ssDNA that recruits RPA. Although the mechanisms leading to the subsequent release of Ku remain obscure, for about 40% of seDSBs induced by the topoisomerase I poison camptothecin (CPT), they involve the coordinated nuclease activities of MRE11 and CtIP, and activation by ATM [33]. Regulation of end resection also involves p53 binding protein 1 (53BP1), effector molecules [34] and the helicase HELB [35]. Recently, several splicing factors have been involved in DNA end resection, including ZNF830 [36], Aquarius [37] and XAB2 [37, 38], by mechanisms that remain to be elucidated.

Targeting DNA repair through inhibition of components of the DNA damage response (DDR) has emerged as an important therapeutic approach against many cancers [39]. As a step to identify novel targets for the sensitization of GBM cells to TMZ, we carried out a shRNA screen for DDR genes that are required for cell proliferation in the presence of TMZ. Here, we report the characterization of XAB2, one of the top hits of this screen, and describe a novel role for XAB2 in promoting Ku eviction and HR at seDSBs.

## Results

### XAB2 is involved in the response of GBM cells to temozolomide

As a step to identify novel TMZ sensitizers in GBM cells, we performed a pooled shRNA screen targeting 574 DDR genes for gene depletions that conferred long-term loss of proliferation in the presence of TMZ (but not vehicle) to the GBM cell line NCH644 (Fig. 1a). The screen was carried out in duplicate, using TMZ at a clinically-achievable concentration of 60 μM under serum-free conditions and in a monolayer format to ensure uniform TMZ distribution. Cells were harvested following 8 and 15 cumulative population doublings for DNA extraction, library preparation and shRNA read counting via high-throughput sequencing (Fig. 1b). Twenty-six genes were identified as prioritized candidates by overlapping the top ranking genes from 3 analysis algorithms, among which *XAB2* (Fig. 1c,d), whose contribution to TMZ-induced DDR was hitherto unexplored.

**Fig. 1.**
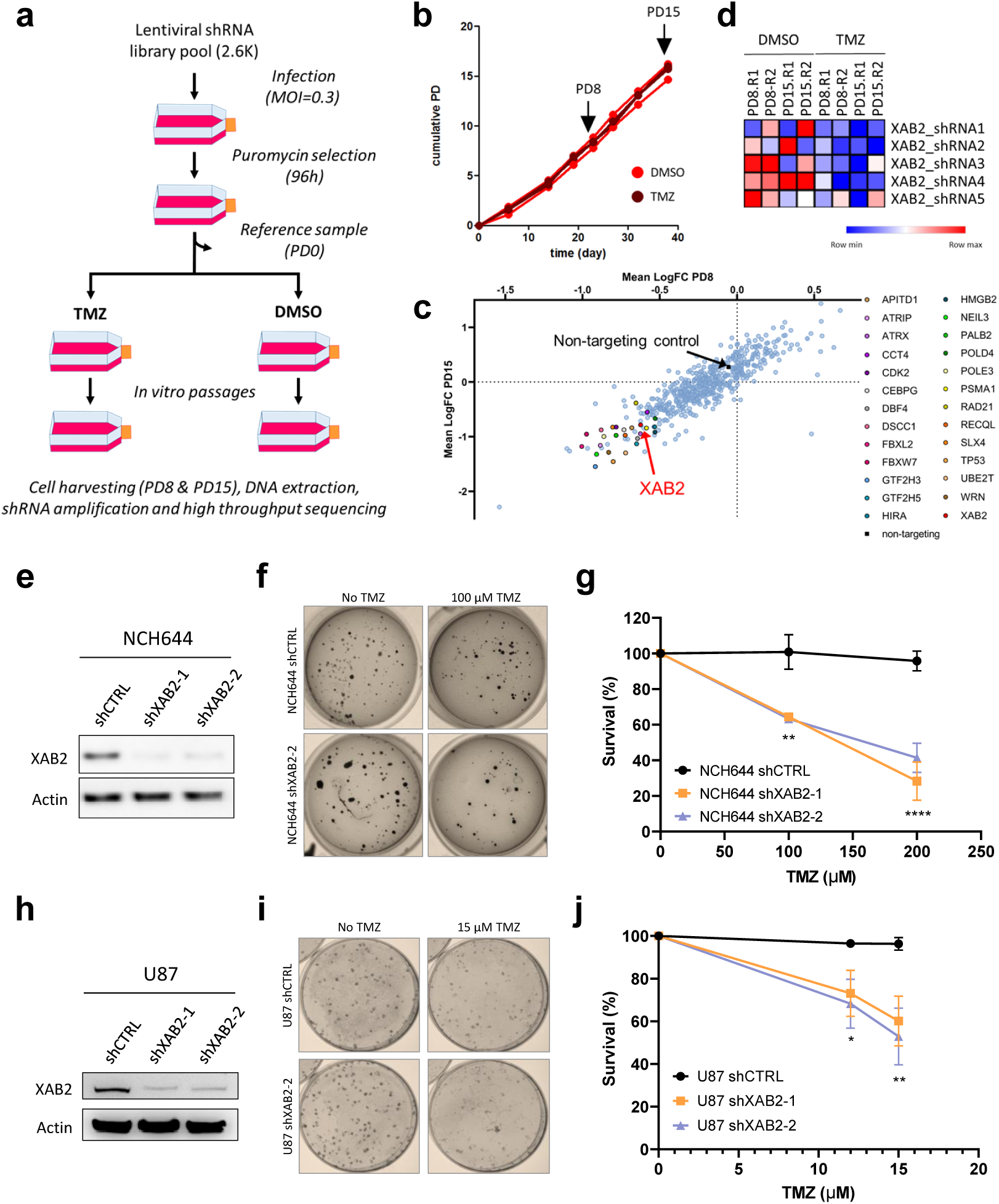
XAB2 promotes resistance to TMZ in GBM cells. **a**, Outline of the shRNA screen. NCH644 cells were infected with a pool of shRNA lentiviral particles targeting 574 gene components of the DNA damage response. Following puromycin selection and harvesting of a reference sample (PD0), cells were split into 2 arms and exposed to 60 μM TMZ or vehicle (DMSO), respectively. Genomic DNA was extracted from the surviving cell populations harvested after 8 and 15 cumulative population doublings (PD). DNA libraries were prepared and sequenced using a MiSeq platform. MOI: multiplicity of infection. **b**, NCH644 cell proliferation assessed under TMZ or DMSO, during the 2 experimental screen replicates. Arrows indicate the PD8 and PD15 time points at which cells were harvested. **c**, Means of the shRNAs fold changes (FC) (TMZ versus DMSO) for each gene of the libray at PD15 compared to PD8. Highlighted are the genes consistently identified as prioritized candidates by overlapping the top ranking genes from 3 analysis algorithms. **d**, Heatmap showing the relative depletion of each of the 5 shRNAs targeting *XAB2* present in the library, in cells treated with TMZ compared to DMSO, at the indicated PD, in the 2 replicates of the screen. R1 and R2: experimental replicate 1 and 2, respectively. Normalized count values are depicted using a red (high) to blue (low) color key. **e**-**j**, XAB2 depletion results in increased sensitivity to TMZ in NCH644 cells (**e**-**g**) and U87 cells (**h**-**j**). **e, h**, representative immunoblots illustrating the efficiency of XAB2 depletion achieved by 2 independent shRNAs (shXAB2-1 and shXAB2-2) as compared to a control, non-silencing shRNA (shCTL). Actin was used as a loading control. **f, i**, Clonogenic survival assays. Cells were exposed to the indicated concentrations of TMZ or vehicle (DMSO) for 2 h and allowed to form colonies before being stained with crystal violet. **g, j**, Quantification of the clonogenic assays following cell counting with ImageJ. Data are the average of n=3 or more biological replicates. Bars represent mean ± s.e.m. Significant differences between specified comparisons were assessed by 2-ways ANOVA, and highlighted by stars (*P<0.05; **P<0.01; ****P<0.0001).

We validated *XAB2* as a novel TMZ sensitizer using clonogenic assays with NCH644 cells (GBM cancer stem-like cell line) (Fig. 1e-g) and U87 cells (GBM adherent cell line) (Fig. 1h-j) expressing a control, non-silencing shRNA (shCTRL) or 2 independent shRNAs targeting XAB2.

### XAB2 is important for the repair of DSBs associated with O^6^-meG lesions left unrepaired by MGMT

Given the proposed role of XAB2 in promoting HR [38], we first examined the impact of XAB2 depletion on the repair of DSBs induced by TMZ. We treated control and XAB2-depleted NCH644 cells with TMZ for 2 h and visualized γH2AX foci, a DSB marker, by indirect immunofluorescence (IF) after a 72 h recovery period in drug-free medium, corresponding to ∼2 cell cycles after DNA damage induction. Compared to control cells, XAB2 depletion led to a ∼2 fold increase in the number of foci observed following exposure to TMZ (Fig. 2a,b), and this accumulation was corroborated by comet assay analysis under neutral conditions, which monitors DSB formation (Fig. 2c,d).

**Fig. 2.**
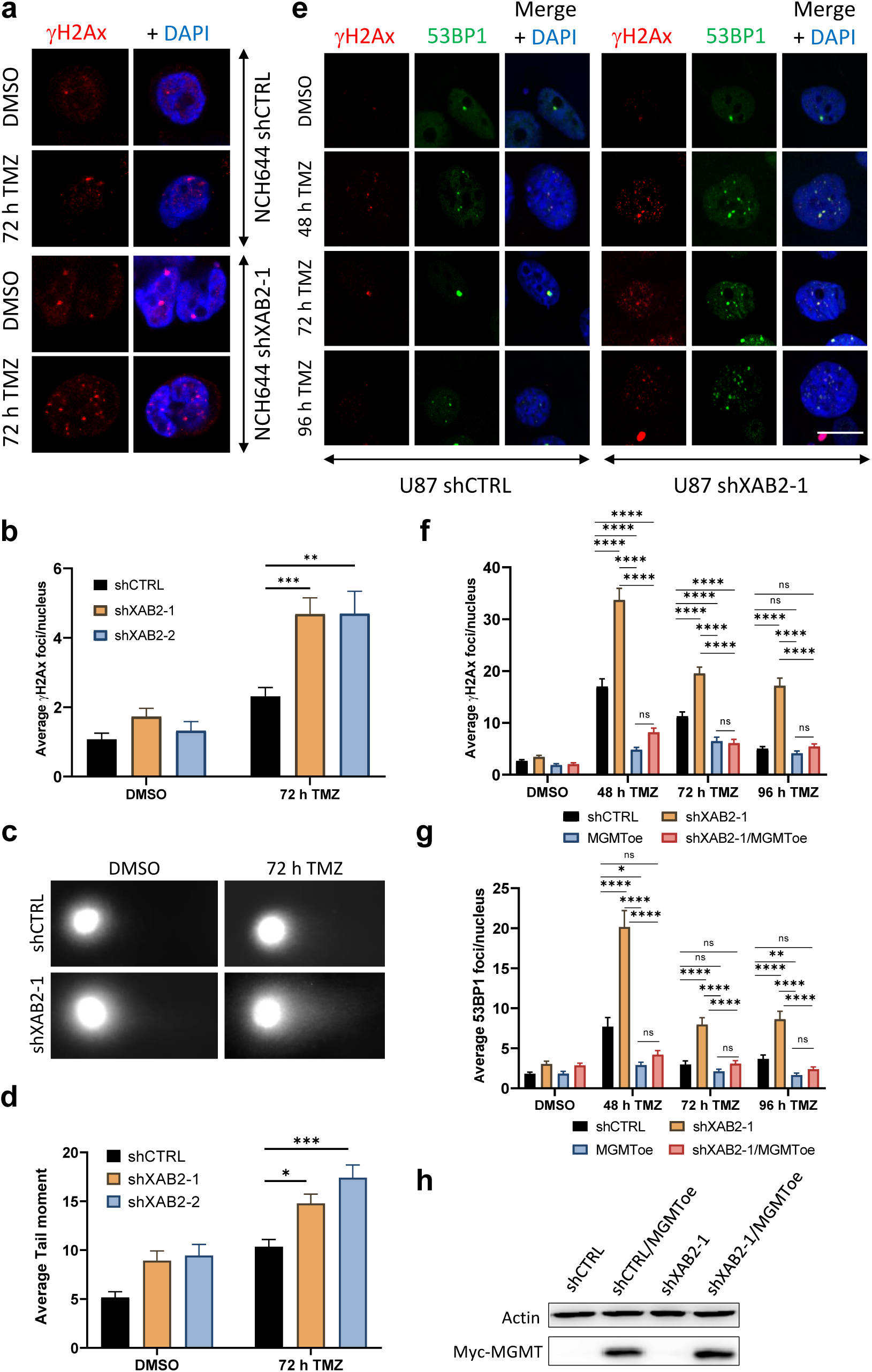
XAB2 depletion impairs the repair of DSBs induced by O6-meG left unrepaired by MGMT. **a**-**b**, Representative immunofluorescence images of γH2AX foci (red) in NCH644 cells expressing the indicated shRNAs, following a 2 h exposure to 100 µM TMZ (or vehicle) and a 72 h recovery period (**a**), and related quantification (**b**). DNA was counterstained with DAPI (blue). **c**-**d**, Control and XAB2-depleted NCH644 cells were treated with 100 µM TMZ (or vehicle) for 2 h and DNA damage was assessed by neutral comet assay after 72 h. Shown are representative images (**c**), and related quantification of the average mean tail moments (30 cell/sample/experiment (n=3) (**d**). **e**, Representative immunofluorescence images used for the quantification of γH2AX (red) and 53BP1 (green) foci in control and XAB2-depleted U87 cells following exposure to 15 µM TMZ (or DMSO) and recovery in drug-free medium for 48, 72 and 96 h. **f**-**g**, Quantification of the average γH2AX foci (**f**) and 53BP1 foci (**g**) per nucleus in control and XAB2-depleted U87 cells as well as in derivative cells ectopically overexpressing MGMT. **h**, Immunoblot analysis of MGMT overexpression in U87 cells. Scale bar= 5µm. The images are representative of 3 or more independent biological repeats. Bars represent mean ± s.e.m. Significant differences between specified comparisons were assessed by Kruskal-Wallis test and are highlighted by stars (*P<0.05; **P<0.01; ***P<0.001; ****P<0.0001).

To obtain more insights into the DSB repair defects associated with XAB2 depletion, we characterized adherent U87 GBM cells which are more amenable to IF microscopy. As observed with NCH644 cells, TMZ exposure led to an increase in γH2AX foci in control U87-shCTRL cells, displaying maximal foci accumulation at 48 h (i.e. about 2 cell cycles) and a return to background levels at the later time points (Fig. 2e,f). TMZ exposure led to a stronger increase in γH2AX foci in XAB2-depleted cells compared to control cells. In addition, unlike for control cells, we observed only a moderate decrease in γH2AX foci at the later time points in XAB2-depleted cells (Fig. 2e,f). Similar observations were made with another DSB marker, 53BP1 (Fig. 2e and 2g). Thus, XAB2 is required for the repair of TMZ-induced DSBs.

TMZ-induced DSBs can arise when the replication fork collides with ssDNA gaps generated during MMR futile cycling at O^6^-meG lesions left unrepaired by MGMT. To explore this question, and since U87 cells do not express MGMT, we examined the impact of XAB2 depletion on TMZ-induced γH2AX and 53BP1 foci formation in otherwise isogenic U87 derivatives ectopically expressing MGMT (Fig. 2h). Stable expression of MGMT prevented the accumulation of γH2AX and 53BP1 foci associated with XAB2 depletion at all time points (Fig. 2f,g). Similarly, ectopic MGMT overexpression prevented TMZ-induced γH2AX foci accumulation in XAB2-depleted NCH644 cells (MGMT-positive) (Supplementary Fig. 1a,b). Taken together, these results indicate that XAB2 promotes the repair of O^6^-meG-associated seDSBs. To gain support for the notion that XAB2 operates at seDSBs resulting from collapsed replication forks, we examined cells treated with hydroxurea (HU), which depletes the dNTP pool and causes fork stalling (upon short exposure) or collapse (upon long exposure) [40]. As expected, short incubation with HU did not cause significant γH2AX foci accumulation in control and XAB2-depleted cells whereas prolonged exposure led to a significant increase in γH2AX and RAD51 recombinase foci accumulation in XAB2-depleted cells compared to control cells (Supplementary Fig. 2a,b).

### XAB2-depletion leads to increased engagement of non-homologous end-joining for the repair of seDSBs during S phase

As HR is the preferred repair pathway for seDSBs resulting from collapsed replication forks and RAD51 recombinase acts upon RPA-coated resected seDSBs, we next examined RPA and RAD51 foci formation in TMZ-treated cells. Control and XAB2-depleted cells displayed a similar increase in RPA foci 48 h after TMZ treatment, indicating efficient end resection (Fig. 3a,b). Moreover, loss of XAB2 led to an increased accumulation of RAD51 foci under these conditions, which paralleled the increase in γH2AX foci already seen at this time point (Fig. 3a-d). Notably, unlike γH2AX and 53BP1 foci, RAD51 foci did not accumulate significantly at the later time points (Fig. 3c,d). As XAB2 depletion did not affect the percentage of cells in S/G2 and G1 phase at the time points considered, as assessed by Cyclin-A staining (Supplementary Fig. 3), these observations suggest that, in the absence of XAB2, RAD51 did not act on a subset of seDSBs induced by TMZ.

**Fig. 3.**
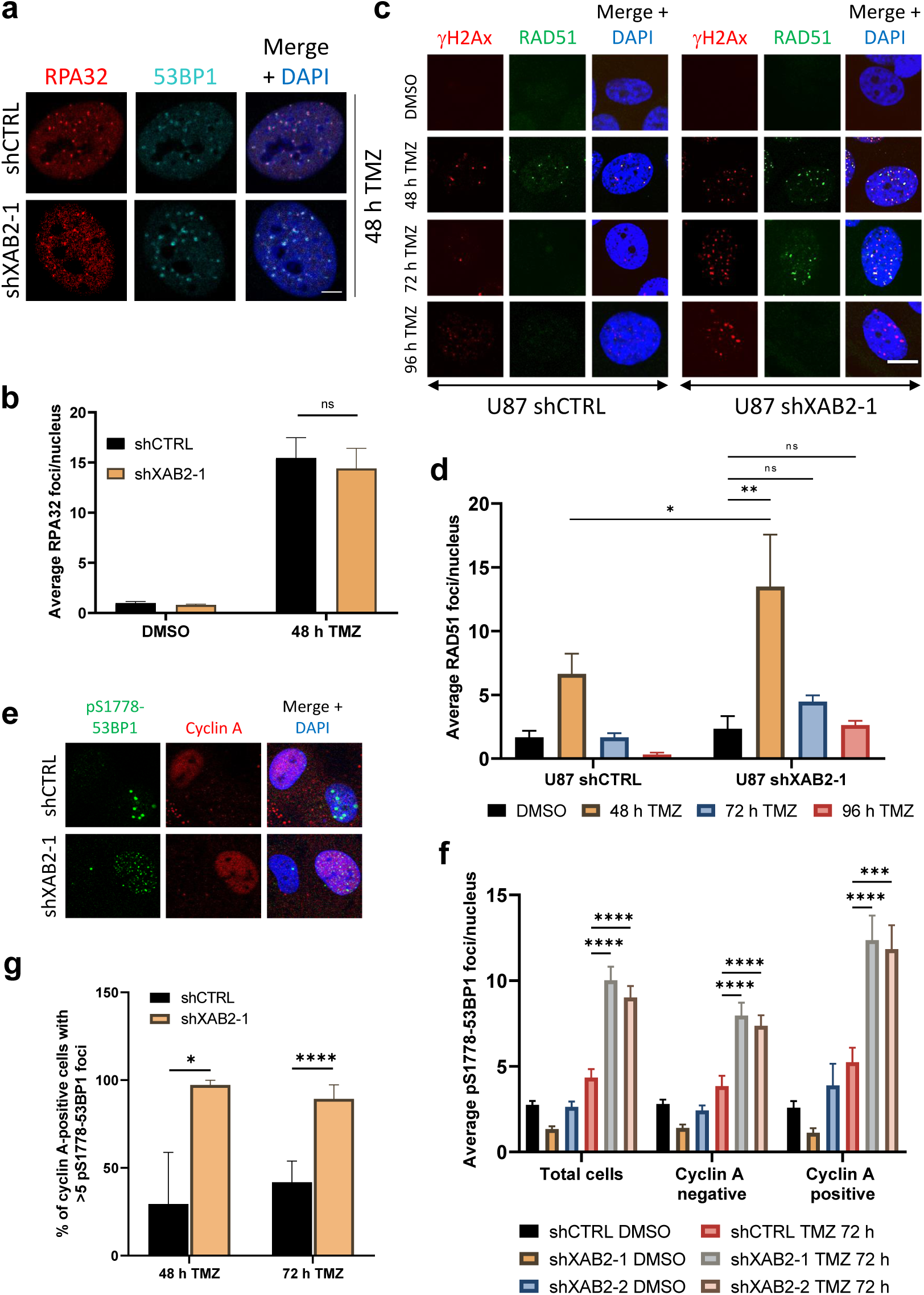
XAB2 depletion leads to the increased engagement of NHEJ in S/G2. **a**-**b**, Representative immunofluorescence images of RPA32 foci (red) and 53BP1 foci (green) in control and XAB2-depleted U87 cells exposed to 15 μM TMZ (or DMSO) for 2 h and allowed to recover in drug-free medium for 48 h (**a**), and related quantification of the average number of RPA32 foci per nucleus (**b**). **c**-**d**, Representative immunofluorescence images of RAD51 foci (green) in control and XAB2-depleted cells exposed to 15 μM TMZ (or DMSO) for 2 h and allowed to recover in drug-free medium for the indicated times (**c**), and related quantification (**d**). Also stained in (**c**) are γH2AX foci (red). **e**-**f**, Representative immunofluorescence images of control and XAB2-depleted U87 cells following exposure to 15 μM TMZ (or DMSO) for 2 h, 72 h recovery in drug-free medium and processing for IF analysis of pS1778-53BP1 foci and Cyclin A (**e**), and related quantification of the average number of pS1778-53BP1 foci per nucleus, in the total cell population, as well as in Cyclin A-negative and -positive cell subpopulations (**f**). **g**, Quantification of the percentage of Cyclin A-positive cells with > 5 pS1778-53BP1 foci per nucleus in control and XAB2-depleted U87 cells exposed to 15 μM TMZ (or DMSO) for 2 h before being allowed to recover in drug-free medium for the indicated times. Scale bar=5µm. The images are representative of 3 or more independent biological repeats. Bars represent mean ± s.e.m. Significant differences between specified comparisons were assessed by Mann-Whitney test in **b** and **g**, 2-ways ANOVA in **d**, Kruskal-Wallis test in **f** and are highlighted by stars (*P<0.05; **P<0.01; ***P<0.001; ****P<0.0001).

We next tested if NHEJ gained more prominence in XAB2-depleted cells. To this end, we examined foci formation by the Ser1778-phosphorylated form of 53BP1 (pS1778-53BP1), which has been implicated in NHEJ [41, 42] and the repair of broken replication forks [43]. To relate our findings to cell cycle stages and distinguish cells in S/G2 from G1 cells, we also visualized Cyclin A. In Cyclin A-negative (G1) cells, XAB2 depletion caused an increase in the number of TMZ-induced pS1778-53BP1 foci observed after 72 h (Fig. 3e,f), consistent with the extra burden of unrepaired damage associated with defective HR. Importantly, under these conditions, XAB2 loss also resulted in a ∼ 2.4 fold increase in pS1778-53BP1 foci in Cyclin A-positive (S/G2) cells (Fig. 3e,f). Furthermore, the percentage of S/G2 cells with more than five pS1778-53BP1 foci reached ∼87% in TMZ-treated XAB2-depleted cells, compared to ∼41% in control cells (Fig. 3g). A comparable increase in NHEJ engagement in S/G2 was already observed at 48 h (Fig. 3g).

One prediction of the increased engagement of NHEJ in the repair of TMZ-induced seDSBs in XAB2-depleted S/G2 cells is that it should be associated with increased genetic instability. Consistent with this notion, loss of XAB2 increased the number of chromosomal aberrations detected in metaphase spreads by 2.5 fold following TMZ exposure (Supplementary Fig. 4a,b).

### Loss of XAB2 is associated with Ku retention on resected DNA repair intermediates and abortive homologous recombination

Although our data indicate that resection and ssDNA/RAD51 associations can occur in the absence of XAB2, the increased engagement of NHEJ observed in S phase in XAB2-depleted cells suggests that loss of XAB2 inhibits HR in some way. Based on previous work [33], we considered the possibility that XAB2 depletion led to the Ku persistence at seDSB termini. Following pre-extraction with Triton X-100 and RNAse A to enable IF analysis of Ku [44], we found that TMZ treatment led to a ∼ 1.5 fold increase in Ku80 foci in XAB2-depleted cells compared to control cells (Fig. 4a,b). ATM defines one pathway for Ku release at seDSBs induced by CPT [33]. In agreement with this study, inhibition of ATM kinase activity using the ATM inhibitor (ATMi) KU-55933 [45] also resulted in increased Ku80 foci formation in TMZ-treated cells (Fig. 4a). Notably, ATM inhibition further increased the number of Ku80 foci elicited by XAB2 depletion (Fig. 4a). Importantly, Ku80 foci detected following exposure to TMZ were associated with resected seDSBs as assessed by the accumulation of RPA foci. Thus, loss of XAB2 led to a ∼3 fold increase in the frequency of Ku80-RPA32 colocalized foci (Fig. 4b-d) compared to control cells. However, contrary to ATM inhibition, which supported end resection whilst preventing RAD51 foci formation in CPT-treated cells [33], XAB2 depletion resulted in a ∼ 2.9 fold increase in the frequency of Ku80-RAD51 foci (Fig. 5e-g). Of note, the colocalization data of Fig. 4c and 4f suggest that XAB2 is required to prevent Ku retention at about a third of seDSB termini.

**Fig. 4.**
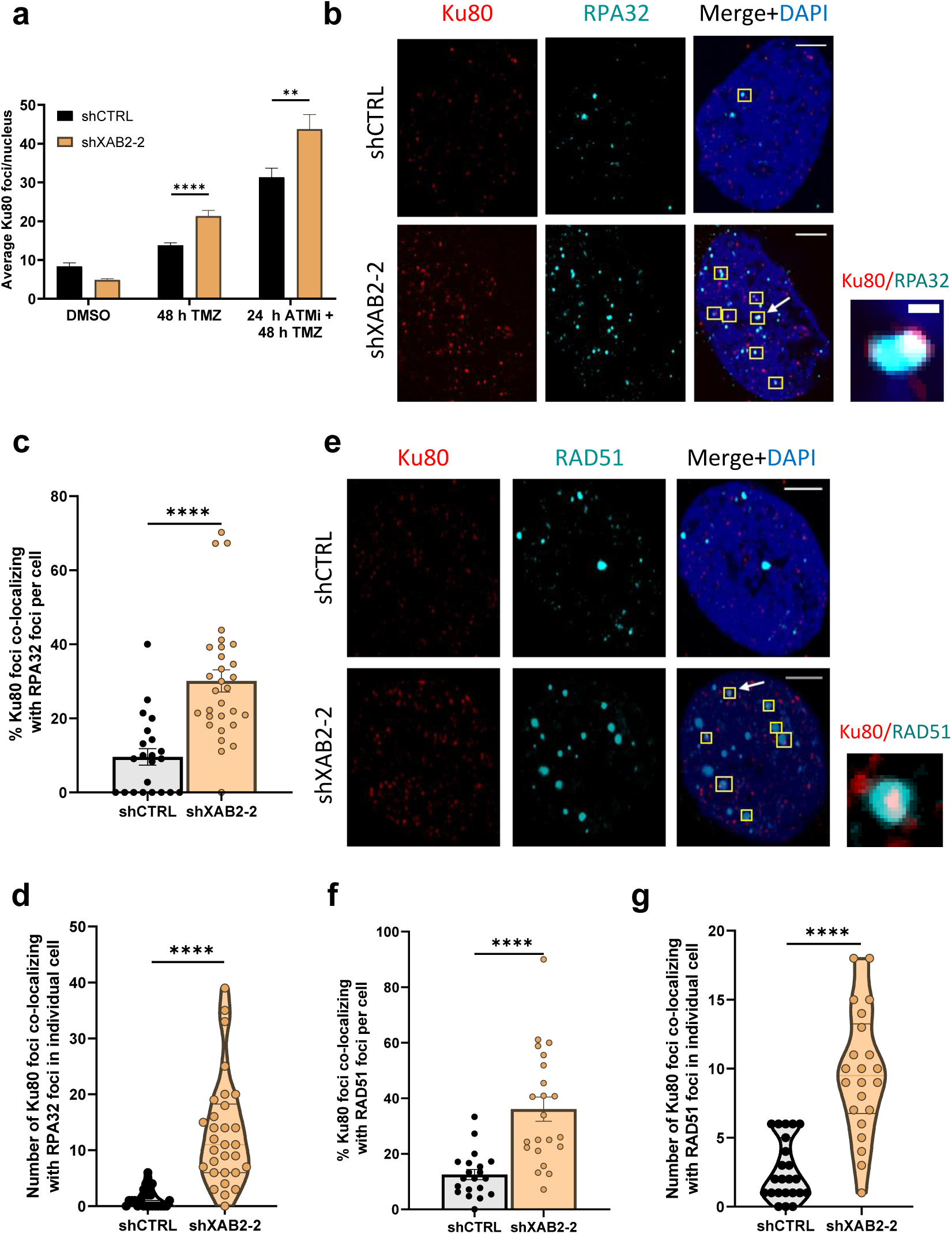
XAB2 prevents Ku retention on seDSBs induced by temozolomide. **a**, Quantification of the average number of Ku80 foci detected by immunofluorescence microscopy in control and XAB2-depleted U87 cells exposed to 15 μM TMZ (or DMSO) for 2 h and allowed to recover for 48 h in the absence or presence of the ATMi KU-55933. **b**, Representative immunofluorescence images used for the quantification of the number of colocalizing Ku80 (red)-RPA32 (cyan) foci following exposure of control and XAB2-depleted U87 cells to TMZ and recovery for 48 h. Examples of colocalized foci are highlighted by yellow squares, with one representative example (indicated by an arrow) shown in the close up section (scale bar = 0.1 μm). **c**-**d**, Percentage of colocalized Ku80-RPA32 foci per cell following exposure to TMZ as in **c**, quantified based on foci examination in individual cells, as presented in a violin plot (**d**). **e**-**g**, Same as **b**-**d** for the analysis of Ku80 (red) and RAD51 (cyan) foci colocalization. Scale bar = 5µm. The images are representative of 3 independent biological repeats. Bars represent mean ± s.e.m. Significant differences between specified comparisons were assessed by a t-test (unpaired, 2-tails) and are highlighted by stars (*P<0.05; **P<0.01; ***P<0.001; ****P<0.0001).

**Fig. 5.**
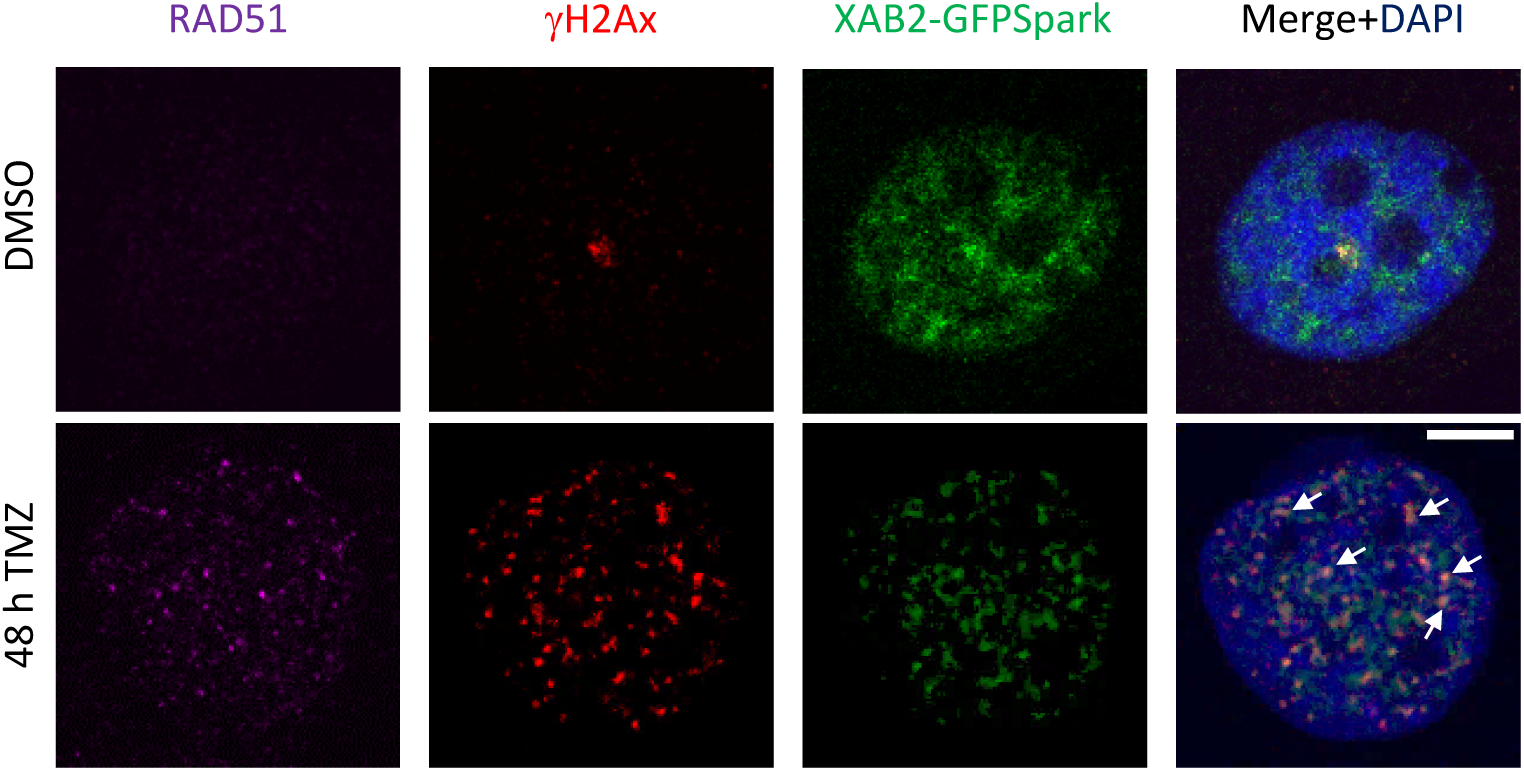
Colocalization of XAB2 with DSBs and RAD51 following DNA damage induced by temozolomide. U87 cells expressing XAB2-GFPSpark were exposed to TMZ for 2 h and allowed to recover for 48 h in drug free media before fixation and IF for RAD51 (magenta), γH2AX (red). Scale bar=5µm. White arrows highlight examples of colocalization between XAB2-GFPSpark, gH2AX and RAD51 foci. The images are representative of 3 independent biological repeats.

We next sought to extend our findings with U87 cells to seDSBs induced by CPT. Exposure to 1 μM CPT for 1 h generated equal levels of DNA damage in control and XAB2-depleted cells, as assessed by γH2AX foci formation (Supplementary Fig. 5a). As seen for TMZ, CPT treatment led to an increase in Ku80 foci in XAB2-depleted cells compared to control cells (Supplementary Fig. 5b,c), which was exacerbated in the presence of ATMi (Supplementary Fig. 5b,c), in agreement with previous work [33]. Furthermore, loss of XAB2 increased the frequency of CPT-induced Ku80-RPA32 colocalized foci (>2.5 fold) and Ku80-RAD51 colocalized foci (>1.7 fold) (Supplementary Fig. 5d-f). Finally, whereas the ATM pathway for Ku eviction involves phosphorylation of CtIP [33], we did not observe significant differences in the levels of TMZ- or CPT-induced hyper-phosphorylation of ATM and CtIP in XAB2-depleted cells compared with control cells (Supplementary Fig. 6a-c). We also verified that loss of XAB2 did not affect ATM activity, as determined by the phosphorylation of the ATM target KAP1 [46] in cells treated with TMZ (Supplementary Fig. 5a). Taken together, our data suggest that ATM and XAB2 identify separate pathways for Ku eviction from seDSB termini, with different impacts on RAD51 filament assembly.

In a previous study investigating the role of XAB2 in HR in the U2OS human osteosarcoma cell line, siRNA-mediated depletion of XAB2 resulted in a decrease in CPT-induced chromatin-bound RPA, as assessed by flow cytometry analysis, which was interpreted as indicative of defective end resection [38]. However, similar to U87 cells, we found that shRNA-mediated XAB2 depletion in U2OS cells did not prevent RPA32 and RAD51 foci formation following exposure to CPT (Supplementary Fig. 7a-f), suggesting that end resection was not impaired in these cells. Ruling out possible effects linked to the RNAi method used to deplete XAB2, targeting XAB2 with specific siRNAs (Supplementary Fig. 8a) did not impact RPA32 foci formation (Supplementary Fig. 8b,c). To complement our analysis of RPA foci, we monitored DNA end resection in the siRNA-treated cells through IF visualization of bromodeoxyuridine (BrdU)-labelled ssDNA. XAB2 depletion did not decrease the number/intensity of BrdU foci elicited by CPT compared to control cells (Supplementary Fig. 8d-g). Finally, we showed that XAB2 depletion did not affect CtIP and ATM phosphorylation in U2OS cells treated with CPT, as previously seen in U87 (Supplementary Fig. 8h-i). Collectively, our observations suggest that XAB2 is not required for seDSB end resection but prevents Ku retention and abortive HR at seDSBs induced by CPT and TMZ.

### XAB2 is recruited to temozolomide-associated γH2AX foci and colocalizes with RAD51

As XAB2 is composed essentially of 15 tetratricopeptide repeat (TPR) domains [47] and since TPRs function as protein interaction modules and multiprotein complex scaffolds [48], we next wanted to obtain evidence for a direct role of XAB2 in HR-mediated seDSB repair. Using a XAB2-GFPSpark fusion expressed in U87 cells, we observed a predominantly diffuse nuclear signal in untreated cells, with granular structures that may represent splicing speckles [49] (Fig. 5). However, TMZ treatment resulted in the accumulation of XAB2-GFPSpark foci colocalizing with γH2AX and/or RAD51 foci (Fig. 5). These observations suggest that the impact of XAB2 depletion on Ku retention at seDSBs stems from a direct role for XAB2 at collapsed replication forks.

### Rescue of the DNA repair deficiencies in XAB2-depleted cells by overexpression of RAD51 and RAD52

Previous studies in yeast have shown that RAD51 and RAD52 overexpression could suppress defective BIR caused e.g., by dysfunctional RPA stabilization of ssDNA [50]. We found that stable ectopic overexpression of RAD51 or RAD52 (Fig. 6a-c) decreased the number of TMZ-induced γH2AX foci associated with XAB2 depletion to near background levels, with overexpressed RAD52 affording the strongest rescue at the later time points (Fig. 6b,c). In line with this observation, analysis of the 72 h time points for Cyclin A and pS1778-53BP1 positivity, revealed that overexpression of RAD52, but not RAD51, reduced NHEJ engagement in S/G2 in XAB2-depleted cells to near background levels (Fig. 6d).

**Fig. 6.**
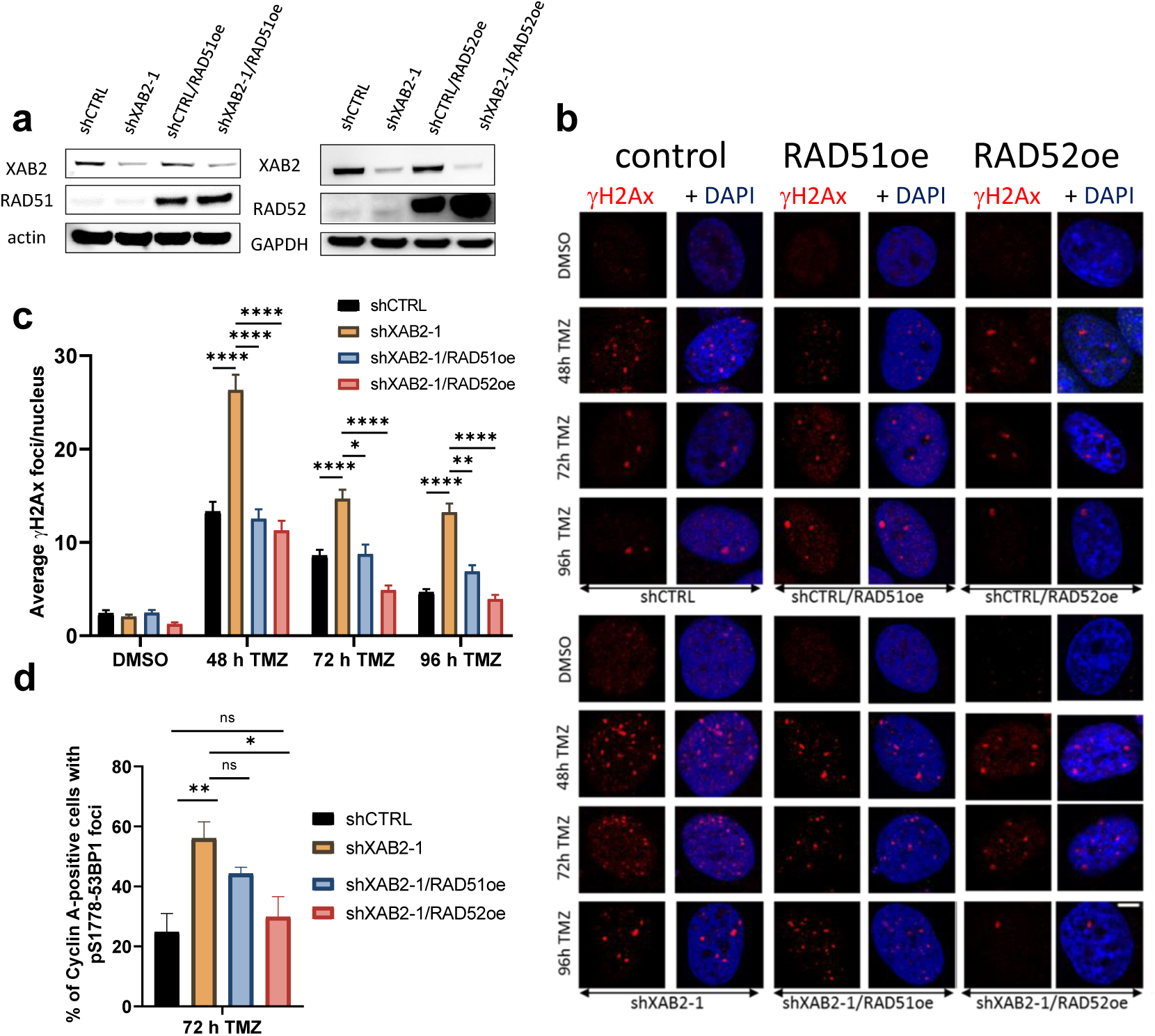
Defective XAB2 is partially rescued by overexpression of RAD51 and RAD52. **a**, Immunoblotting analysis of RAD52 (upper panel) and RAD51 (lower panel) overexpression in control and XAB2-depleted U87 cells. **b**-**c**, Representative micrographs of γH2AX foci (red) detected by immunofluorescence in the indicated cells exposed to 15 μM TMZ (or DMSO) for 2h and allowed to recover in drug-free medium for the indicated times (**b**) and related quantification in (**c**). **d**, Percentage of Cyclin A-positive cells displaying pS1778-53BP1 foci among control and XAB2-depleted cells harboring the indicated constructs, following exposure to 15 μM TMZ (or DMSO) for 2 h and recovery in drug-free medium for the indicated times. Data are the average of n=2 or more biological replicates (30-50 cells/sample/experiment). Scale bar=5µm. The images are representative of 3 independent biological repeats. Bars represent mean ± s.e.m. Significant differences between specified comparisons were assessed by Kruskal-Wallis test in **c** and 2-ways ANOVA in **d** and are highlighted by stars (*P<0.05; **P<0.01; ***P<0.001; ****P<0.0001) (ns = not significant).

### RAD52 inhibition is synthetically lethal with XAB2 depletion

Although we did not identify RAD52 in our shRNA screen, the significant rescue afforded by its overexpression in XAB2-depleted cells prompted us to explore its importance in the response of GBM cells to TMZ. shRNA-mediated RAD52 depletion in U87 cells led to a strong sensitivity to TMZ (Fig. 7a-c). We next tried to generate cells harbouring the dual depletion of XAB2 and RAD52 using RNA interference. In NCH644, no viable double knockdown cells could be obtained (Fig. 7d), suggesting synthetic lethality. In U87, we obtained double knockdown cells, but their clonogenic survival was severely impaired (Fig. 7e,f).

**Fig. 7.**
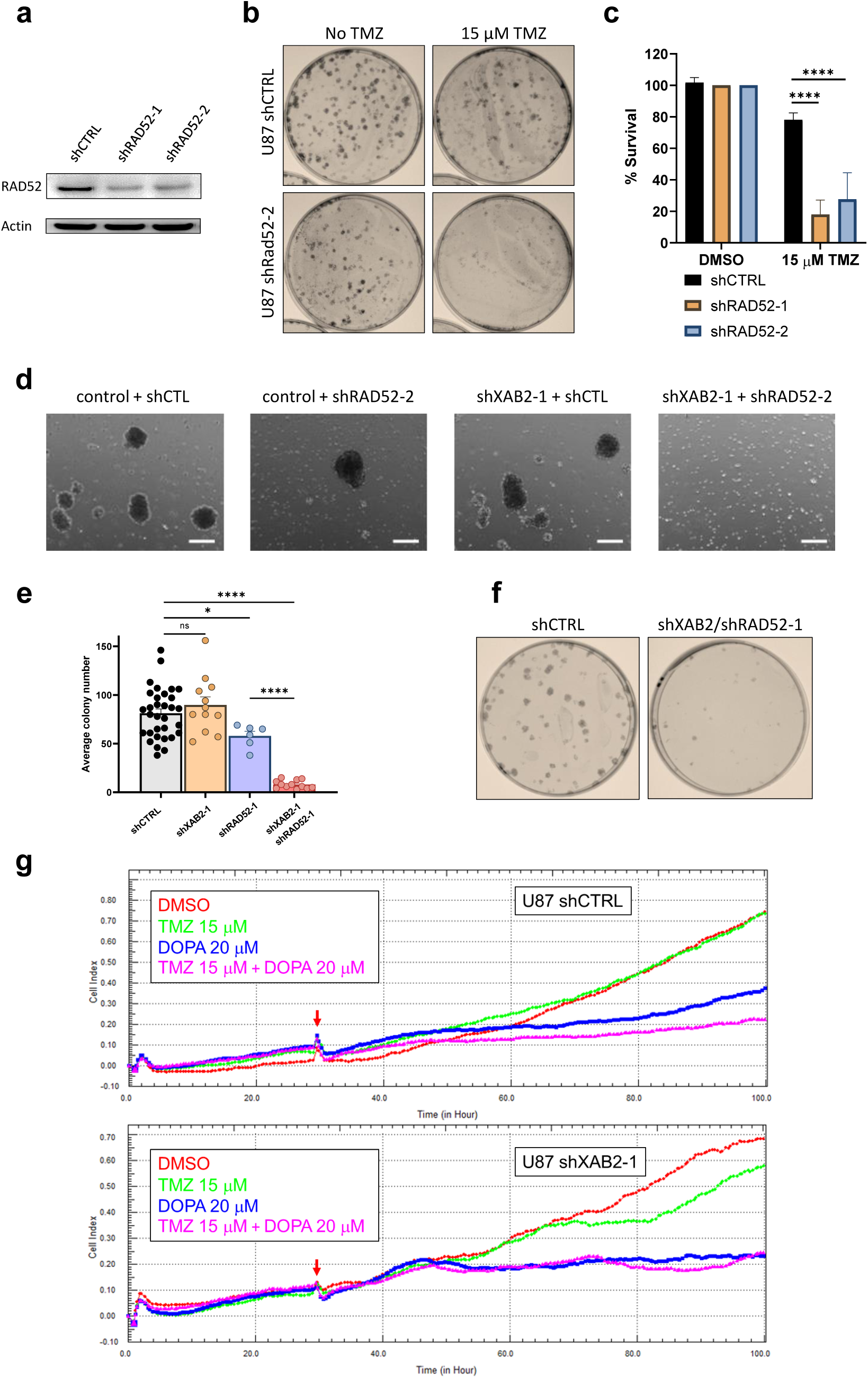
XAB2 shows synthetic lethality with RAD52. **a**, Immunoblots illustrating the efficiency of RAD52 depletion achieved by 2 independent shRNAs (shRAD52-1 and shRAD52-2) as compared to shCTRL. Actin was used as a loading control. **b**-**c**, Clonogenic survival assays carried out with control and RAD52-depleted cells exposed to 15 μM TMZ (or DMSO) for 2 h (**b**) and related quantification (**c**). Cell viabilities are expressed as % relative to that of untreated cells (set at 100%). **d**, Micrographs of control or XAB2-depleted NCH644 cells transduced with either shRAD52 or control (shCTL) shRNAs, taken at day 18 following transduction and selection in puromycin/G418. Control and XAB2-depleted cells, obtained using shRNA constructs expressing the puromycine-resistance marker, were transduced with pLKO-based constructs expressing shCTL or shRAD52-2 shRNAs, followed by additional selection with G418. No spheroid growth was observed in the double knockdown cells. **e**-**f**, Quantification of the clonogenic survival of U87 cells following single or double knockdown of XAB2 and RAD52 (**e**) and representative illlustration of the plating efficiency of the double knockdown cells compared to control cells (**f**). See Fig. 1i and Fig. 7b for illustrations of the single knockdowns. **g**, xCELLigence real-time cell proliferation analysis with control and XAB2-depleted cells exposed to the indicated concentrations of TMZ and/or L-DOPA, or vehicle. Red arrows indicate the time of drug addition. Images shown are the representative of 3 independent experiments.Scale bar = 250 µm Data are the average 3 or more biological replicates. Bars represent mean ± s.e.m. Significant differences between specified comparisons were assessed by one- or two-ways ANOVA and are highlighted by stars (*P<0.05; ***P<0.001).

To corroborate these observations, we carried out real-time cell proliferation analyses following RAD52 inhibition with 6-hydroxy-DL-DOPA (L-DOPA) [51]. In wild-type cells, TMZ had little or no impact on proliferation (Fig. 7g), but exposure to L-DOPA alone impaired proliferation and its effect was exacerbated in combination with TMZ (Fig. 7g), consistent with the impaired clonogenicity of RAD52-depleted cells (Fig. 7b and 7e). As expected, TMZ impaired the proliferation of XAB2-depleted cells. In addition, RAD52 inhibition by L-DOPA abolished the proliferation of XAB2-depleted cells even in the absence of TMZ (Fig. 7b and 7e).

## DISCUSSION

We found that XAB2 defined a novel mechanism to counteract Ku accumulation at seDSBs. Previous studies have shown that an ATM kinase-dependent pathway involving CtIP and MRE11 operates to counteract Ku retention at about 40% of seDSB termini [33]. The synergistic impact exerted by ATM inhibition and XAB2 depletion on Ku retention suggests that the XAB2-dependent pathway operates in parallel to the ATM pathway. The XAB2-dependent pathway differs from the ATM-dependent one in that loss of XAB2 does not prevent RAD51 loading on resected DNA ends. This observation suggests that under certain instances RPA-RAD51 exchange on ssDNA may not require coupling to Ku removal. However, overexpression of RAD51 rescued the DNA damage defect associated with XAB2 depletion, indicating that the ssDNA-RAD51 associations that take place on Ku-bound, resected seDSBs in the absence of XAB2 are not sufficient for efficient recombinational repair (Fig. 8). As XAB2 is present at seDSBs, we propose that XAB2 acts via a direct mechanism to promote Ku eviction from resected seDSBs. The molecular details of this novel pathway and its interplay with the ATM-dependent pathway in counteracting Ku accumulation merit further exploration.

**Fig. 8.**
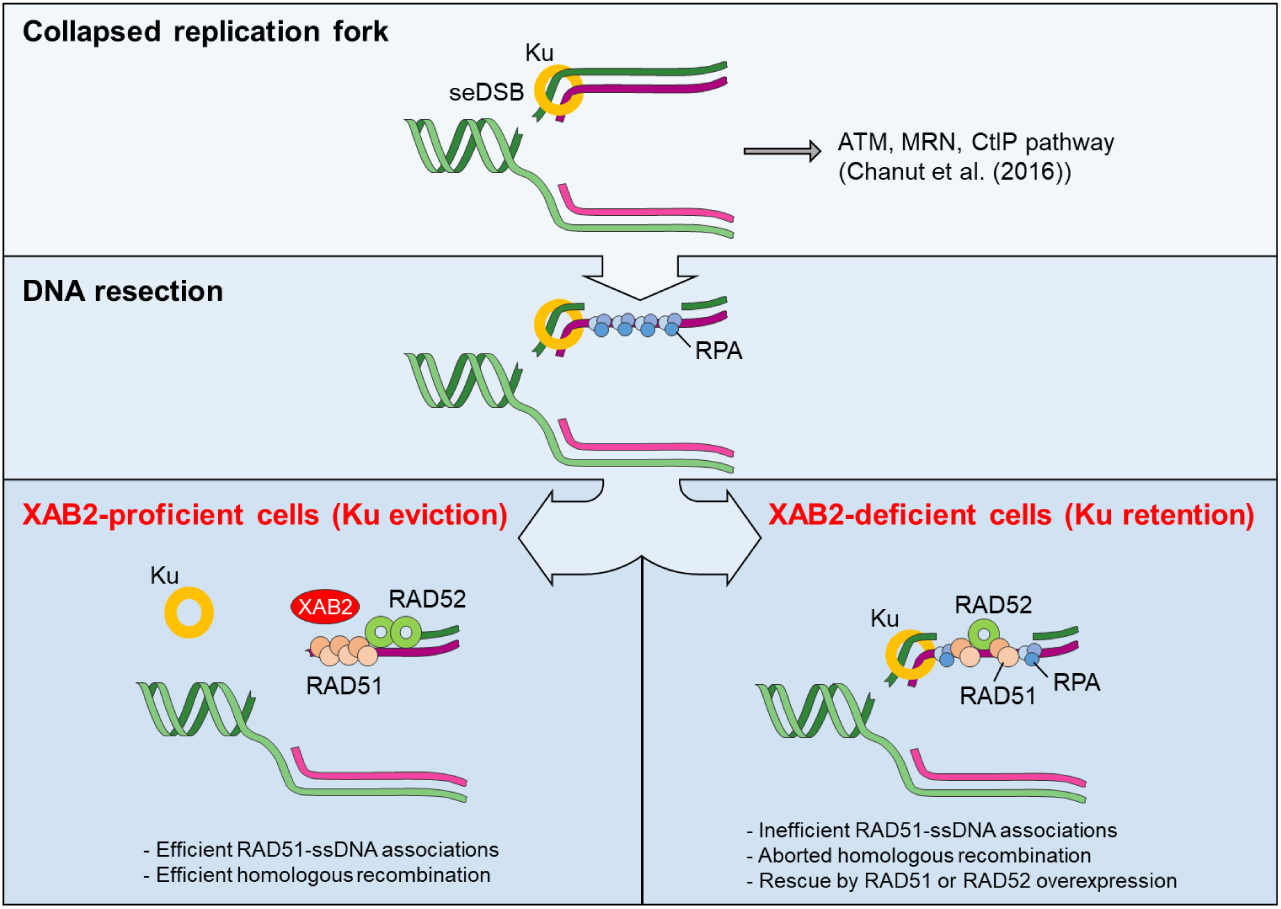
XAB2 promotes Ku eviction from seDSB termini and HR. Proposed model integrating the present findings. XAB2 defines a novel pathway for Ku eviction from a subset of seDSBs resulting from collapsed replication forks. This pathway operates in parallel to an ATM/MRN/CtIP-dependent pathway previously described (ref 33). Whereas the ATM/MRN/CtIP-dependent pathway is critical for RAD51 loading, loss of XAB2 does not prevent interaction between RAD51 and ssDNA generated through end resection. However, the resulting associations are unproductive, leading to increased NHEJ engagement in S/G2. Such defects can be rescued by overexpression of RAD51 or RAD52.

RAD52 overexpression afforded robust rescue of the defects associated with XAB2 depletion. RAD52 can exploit microhomologies [52-54] and its transactions with DNA require moderate resection compared to RAD51-mediated filament formation. Furthermore, RAD52 competes with Ku for binding to DSB free ends generated in the switch region during antibody class-switch DNA recombination [53], suggesting the possibility that, unlike RAD51, it may be able to operate at seDSBs despite the persistence of Ku or that alternatively, it displaces Ku when overexpressed. Future study will be required to test these hypotheses.

RAD52 acts as mediator of ssDNA/RAD51 filaments during seDSB repair by HR in human cells, a role where it can be replaced by BRCA2 [14]. The severe TMZ sensitivity induced by loss of RAD52 and its strong ability to rescue the defective phenotypes associated with XAB2 depletion suggest that RAD52 also exerts a function independently of RAD51, in line with a previous report [15]. Supporting this notion, we found that not all DSBs seen in XAB2-depleted cells are substrates for RAD51. RAD52 mediates MiDAS independently of RAD51 [16]. It is therefore possible that the residual DSBs observed at 72 h and 96 h in XAB2-depleted cells exposed to TMZ, which RAD52 overexpression fully suppressed, reflect in part the participation of XAB2 in MiDAS.

RAD52 inactivation induces synthetic lethality in cells harbouring deficiencies in key HR factors [55-58]. Our observation that XAB2 limits the engagement of NHEJ associated with genome instability can explain why XAB2 was identified in RNAi screens for determinants of PARP inhibitor sensitivity [59, 60] or genes that promote genome stability [61]. Our data suggest that strategies targeting RAD52 may help improve the therapeutic outcome of patients with glioblastoma.

## Methods

### Cell lines, cell cultures and treatments

NCH644 cells were kindly provided by Dr Christel Herold-Mende (Department of Neurosurgery, University of Heidelberg) [62]. NCH644 cells were cultured in in Neurobasal medium (Life Technologies, 21103049) supplemented with 1x B-27 (Life Technologies, 12587010) 2 mM L-Glutamine, 20 U/ml Pen-Strep, 1 U/ml Heparin (Sigma, H3149-25KU), 20 ng/ml bFGF (Miltenyi, 130-093-841) and 20 ng/ml EGF (Provitro, 1325950500). For the shRNA screen, 2D cultures of NCH644 were carried out in the same medium using laminin-coated plates. U87 and U2OS cells were cultured in Dulbecco’s modified Eagle’s medium (Westburg, LO BE12-614F), supplemented with 10% fetal bovine serum (FBS) (Gibco, 10500-064), 50 U/ml Pen-Strep. Cells were routinely subjected to mycoplasma testing using the Mycoplasma PCR ELISA kit (Sigma Aldrich, 11663925910) and tested negative.

Transductants and transfectants were selected using G418 (300 μg/ml) and/or puromycin (1 μg/ml). Stock solutions of Temozolomide (TMZ, 100 mM) (Sigma Aldrich) and Camptothecin (CPT, 50 mM) (Sigma Aldrich) were prepared in DMSO. RAD52 was inhibited using 6-hydroxy-DL-dopa (L-DOPA, Sigma Aldrich) [51].

### shRNA screen, RNA interference and plasmids

The shRNA screen was performed as follows: NCH644 cells grown on laminin-coated plated were infected at an MOI of 0.3 with a 2.6 K custom lentiviral shRNA library constructed in pGIPZ puro vector and targeting 574 DNA Damage Response (DDR) genes (i.e., ∼4.5 shRNAs per gene on average, 700-fold representation)(Thermo Fisher Scientific). Cells were exposed to 1 μg/ml puromycin for 96 h to allow selection of transductants. Following harvesting of a reference sample (PD0), cells were split in 2 arms and cultured under vehicle (DMSO) or 60 µM TMZ (which corresponds to the IC20 for NCH644 interpolated from 2D cytotoxic assays and represents a sub-lethal, therapeutically-reachable dose range [63]). Cells were harvested after cumulative population doubling (PD) 8 and 15 for DNA extraction, library preparation and shRNA read counting via sequencing on an Illumina MiSeq platform. The screen was carried out in duplicate. Screen analysis was performed by comparing shRNAs counts between the vehicle (DMSO) and TMZ treatment conditions using MAGeCK-RRA [64], ScreenBEAM [65] and HitSelect [66]. For each PD, we crossed the lists of the top-15% candidate genes independently identified by the 3 algorithms and kept their intersection. We then considered as prioritized hits a subset of 26 genes consistently appearing in the 3 analyses both at PD8 and PD15 in both replicates.

shRNA-mediated depletion of XAB2 and RAD52 was carried out using pGIPZ (Dharmacon) or pLKO [67] lentiviral vector-based shRNAs. The following pGIPZ-based shRNAs were used: shXAB2-1 (V2LHS_50670; mature antisense: TTGACAGAGAATTGGTTCC), shXAB2-2 (V3LHS_645634; mature antisense: ACAAACGTAGCTGTATTGG), shRAD52-2 (V3LHS_376617; mature antisense: TCATGATATGAACCATCCT), shRAD52-3 (V2LHS_171209; mature antisense: ATTGCTTGAGGGCAAGGAG). Lentiviral vectors expressing non-silencing shRNAs in pGIPZ (puromycine resistance marker) (RHS4346) or in pLKO (neomycin resistance marker) [68] were used as negative controls.

siRNA-mediated depletion of XAB2 was achieved using SMARTpool siGENOME XAB2 siRNAs (Dharmacon), with siGENOME Non-targeting control siRNA pool I used as a non-silencing siRNA control (Dharmacon). siRNA transfection were carried out using Lipofectamine RNAiMAX transfection reagent, as detailed in [38].

Overexpression of MGMT, RAD51 and RAD52: A Myc-tagged MGMT fragment containing 5’-XbaI extremities and 3’-BamHI extremities was generated by PCR amplification and restriction enzyme digestion and cloned into the lentiviral vector pCDH-EF1α-MCS-IRES-Neo and pCDH-EF1α-MCS-IRES-Puro (System Biosciences) pre-digested with the same restriction enzymes. Lentiviral vectors expressing RAD51 and RAD52 under the EF1α promoter were constructed by subcloning of a NcoI(blunt)-BamHI fragment containing RAD51 from pFB530 [69] or a NdeI(blunt)-BamH1 fragment containing RAD52 from pFB581 [70] (kindly provided by Dr. S.C. West, Cancer Research UK) into the lentiviral vectors pCDH-EF1α-MCS-IRES-Neo and pCDH-EF1α-MCS-IRES-Puro (System Biosciences) cut by EcoRI(blunt)-BamHI. All constructs were verified by sequencing.

Plasmid pCMV3-C expressing XAB2-GFPSpark was obtained from Sino Biological. The fusion construct was subcloned into pCDH-EF1α-MCS-IRES-Puro for long-term expression.

### Immunofluorescence (IF) analysis

Cells grown on glass coverslips were incubated with TMZ (or DMSO) for 2 h at 37°C, washed once with media and left to recover in drug-free media for the indicated periods. Cells were then fixed with 4% paraformaldehyde in PBS for 10 min at room temperature (RT), permeabilized for 10 min with 0.5% triton X-100 in PBS and incubated for 30 min at RT in PBS with 2% bovine serum albumin (BSA) to block nonspecific binding. Thereafter the cells were incubated with primary antibodies at 4°C (90 min or overnight, depending on the antibody), washed 3 times with PBS and then incubated with secondary antibodies for 1 h at RT. Cells were counterstained with DAPI and visualized using a Zeiss LSM880 confocal microscope.

For the visualization of Ku foci, cells exposed to DNA damaging agents or vehicle were subjected to pre-extraction using cytoskeleton (CSK) buffer containing 0.7% Triton X-100 and 0.3 mg/ml RNase A (CSK+R) as described [44]. Cells were then fixed with 4% paraformaldehyde in PBS for 15 min at RT, blocked in PBS containing 10% fetal bovine serum and 1% BSA and then incubated with the primary antibodies for 90 min at RT in PBS containing 10% fetal bovine serum and 1% BSA. Following 3 washes with PBS, cells were incubated with the secondary antibodies for 60 min at RT in PBS containing 10% fetal bovine serum and 1% BSA, followed by washes and DAPI counterstaining. The ATMi KU-55933 (Tocris Biosciences) was used as described in experiments with CPT [44]; it was added during the last half of the 48 h recovery period following exposure to TMZ.

Analysis of DNA end resection using immunofluorescence visualization BrdU-labelled ssDNA was carried out as previously described [71] with small modifications. In brief, cells were pre-incubated in the presence of 10 μM BrdU (Sigma) for 24 hours before the treatment with 1 μM CPT for one hour followed by a release of one hour. Cells were subjected to *in situ* fractionation on ice for 10 min using sequential extraction with two different buffers. Pre-extraction buffer 1 (10 mM PIPES, pH 7.0, 300 mM sucrose, 100 mM NaCl, 3 mM MgCl_2_, 1mM EGTA and 0.5% Triton-X100) and followed by pre-extraction buffer 2 (10 mM Tris pH 7.5, 10 mM NaCl, 3 mM MgCl_2,_ 1% Tween20 and 0.5% sodium deoxycholate). Cells were washed three times with PBS followed by fixation with 4% paraformaldehyde (w/v) for 15 min at room temperature. Cells were then fixed for 5 minutes with methanol at -20°C. Cells were washed with PBS, permeabilized in 0.5% Triton X-100 in PBS for 10 min and incubated with blocking buffer (PBS + 5% BSA) for one hour. Cells were incubated overnight at 4°C with anti-BrdU and anti-PCNA antibodies in blocking buffer. Unbound primary antibody was removed by washing in PBS at room temperature followed by incubation with the secondary antibodies in PBS + 1% BSA for one hour at room temperature followed by washes and DAPI counterstaining. Coverslips were mounted onto slides with ProLong Gold antifade mountant. BrdU foci were visualized on a DMI6000B microscope.

### Metaphase spread preparation and analysis

U87 cells (500,000 cell) exposed to TMZ or vehicle as before were allowed to recover for 45 h before being incubated in the presence of 0.5 µg/ml colcemid for a further 3 h at 37°C. Following trypsination and harvesting, cells were then swelled in 5 ml of pre-warmed hypotonic solution (0.54% KCL) added dropwise and incubated at 37°C for 30 min. Thereafter, 5 ml of freshly made, ice-cold fixing solution (ethanol:acetic acid (3:1, v/v)) were added and the preparations were stored overnight at 4°C. Metaphase preparations were dropped onto wet slides, air dried and stained with DAPI for microscopic visualization.

### Neutral Comet assay

Cell were embedded in 0.6% low melting agarose (LMA) and layered onto 0.6% normal agarose pre-coated frosted slides. Slides were then immersed in pre-chilled lysis buffer (2.5 M NaCl, 10 mM Tris-HCl, 100 mM EDTA pH8.0, 0.5% triton X-100, 3% DMSO, pH9.5) for 1.5 h and then washed twice with pre-chilled distilled water (2×10min) and twice with pre-chilled electrophoresis buffer (300 mM sodium acetate, 100 mM Tris-HCl, 1% DMSO, pH8.3) (2×10 min). Slides were equilibrated in fresh electrophoresis buffer for 1 h and then subjected to electrophoresis at 25 V (0.6 V/cm) for 1 h at 4°C. DNA was stained with DAPI, imaged with Zeiss LSM 510 confocal laser scanning microscope and pictures were quantified using Perceptive Instruments Comet Assay IV.

### Protein extracts and western blot analysis

Cells were harvested, washed once with ice-cold PBS and incubated in 1 x RIPA buffer (Millipore, 20-188) supplemented with protease inhibitor cocktail (Roche, 11697498001) and phosphatase inhibitor cocktail (Roche, 04906837001) for 15 min at 4°C. Following centrifugation (16.000 g, 15 min) at 4°C, the lysates were stored at -20°C.

Protein extracts were quantified using the BRADFORD-solution (Bio-RAD, 5000006) and heated for 5-10 min at 95°C in LDS-sample loading buffer (Life Technologies, NP0008) containing 50 mM dithiothreitol (Amersham Biosciences, ref.17-1318-02) before being subjected to SDS-PAGE gel electrophoresis (NuPage^™^ 4-12% Bis-Tris Gel Invitrogen, NP0322box) and semi-dry transfer to a nitrocellulose membrane (iBlot^®^2NC Mini Stacks Invitrogen, IB23002). The membrane was blocked in PBS containing 0.05% tween-20 (PBST) and 5% dry milk for 1 h at RT and incubated overnight with the appropriate primary antibody. Following 3 washes over 10 min with PBST the membrane was then incubated with the appropriate horseradish peroxidase (HRP)-conjugated secondary antibody in PBST containing 5% dry milk for 1-2 h at RT. Following 3 washes over 10 min with PBST at RT, signals were detected using ImageQuant LAS400 (General Electric) and images were captured using a SuperSignal™ West Pico Plus Chemiluminescent Substrate (Thermo Scientific) or SuperSignal™ West Femto Maximum Sensitivity Substrate (Thermo Scientific).

### Colony formation assays

Colony formation assays using U87 cells were carried out as follows: 300 cells were seeded onto 10 cm dishes and treated with TMZ for 2 h at 37 °C in a humidified atmosphere of 5% CO_2_. The medium was then replaced and cells were allowed to form colonies over a period of 15 days. Colonies were stained using crystal violet solution (0.1% Brilliant blue R (Sigma-Aldrich), in PBS) and quantified using ImageJ/FIJI analysis software.

For NCH644 cells, soft agar colony formation assays were carried out as follows: 6×10^3^ NCH644 cells were suspended in 0.3% low melting agarose (LMA) containing 1 ml Neural Stem Cell media (NSC-media, Gibco) supplemented with TMZ or vehicle, and then seeded on top of pre-coated 1 ml 0.6% LMA containing NSC-media in the 6well plate. Cells were incubated for 21 days at 37°C with twice-weekly fresh medium supplementation (200 µl). After 3weeks the cells plates were stained with crystal violet for imaging with ImageQuant TL and colony quantification.

### Cell proliferation assays

NCH644 cells were seeded medium containing TMZ or vehicle (0.6×10^6^ cells in 5 ml). They were then split every 3-4 days under subconfluent conditions while remaining under TMZ or DMSO treatment. At each time point, the cumulative population doublings (PD) was calculated as follows:

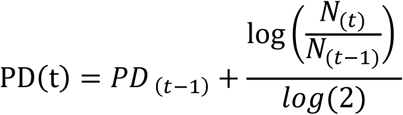

where N(t) is the number of cells counted at time (t) and N(t-1) is the number of cells seeded at the previous time point, (t-1). Cumulative population doublings were plotted against time.

Real-time cell proliferation analyses with U87 cells were carried using a xCELLigence Real-Time Cell Analyis (RTCA) instrument (ACEA Biosciences Inc.) in a 96-well format.

### Primary and secondary antibodies used for IF

Primary antibodies: XAB2 (abcam, ab129487, dilution 1:1000), RAD51 (Merck, Ab-1, PC130, dilution 1:1000), γH2AX (Millipore, Cat. No. 05-636, Clone JBW301, dilution 1:1000), 53BP1 (abcam, ab36823, dilution 1:1000), pS1778-53BP1 (Cell signalling, 26755, dilution 1:1000), Phospho RPA32 (S4/S8) (Bethyl Laboratories, A300-245A, dilution 1:1000), Ku80 (abcam, ab80592, dilution1:1000), BrdU (GE healthcare, RPN202, dilution 1:1000), PCNA (Chromotek, 16D10-100, 1:500). Secondary antibodies: Alexa Flour 647 (Invitrogen, A-21235, dilution 1:500), Alexa Flour 555 (Invitrogen, A-21422, dilution 1:500), Alexa Flour 488 (Invitrogen, A-11001, dilution 1:500), Alexa Fluor 568 (Invitrogen, A-11077, dilution 1:500).

### Primary and secondary antibodies used for western blotting

Primary antibodies: RAD51 (Merk, Ab-1, PC130, dilution 1:1000), RAD52 (Thermo Fisher, PA5-65036, dilution 1:500), XAB2 (abcam, ab129487, dilution 1:1000), CtIP (Active Motif, 61141, dilution 1:500), ATM (abcam, ab32420 (Y170), dilution: 1:1000), P-ATM (abcam, ab81292 (pS1981), dilution 1:50000), P-KAP1 (Bethyl, IHC-00073 (pS824) dilution 1:200). Secondary antibodies: HRP rabbit (Jackson Laboratory, Cat no 111-035-003, dilution 1:50,000), HRP mouse (Amersham/Sigma, Cat no GENA931-1ML, dilution 1:10,000), HRP mouse (Santa Cruz, sc-516102, dilution 1:10,000).

### Statistics and reproducibility

All the error bars are the standard error of mean (s.e.m.), unless mentioned otherwise in the legend. All the graphs are derived from two to five independent repeats. According to the number of samples, statistical significance (P values) of the difference between the means was determined by student *t-test* (two-tailed, upaired), one- or two-ways ANOVA. In the case of non-normally distributed datasets, non-parametric statistical tests have been preferred: Mann-Whitney test for comparisons between two samples and Kruskal-Wallis test for multiple comparisons. Statistical significance was always denoted as follow: ns=not significant; \**p*<0.05; \*\**p*<0.01; \*\*\**p*<0.001; \*\*\*\**p*<0.0001. All statistical analysis was perfprmed using GraphPad Prism 8 (GraphPad Software).

## Ackowledgements

This work was supported by Télévie/Fonds National de la Recherche (F.R.S.-FNRS)/Fonds National de la Recherche du Luxembourg (FNR) (grant 7.4503.11 to H.E. and E.V.D.; grant 7.4633.16 to A.B.S and E.V.D), the Doctoral School for Systems and Molecular Biomedicine, University of Luxembourg (Grant to H.E.), FNR (PRIDE grant to L.P.), and a Canadian Institutes of Health Research Foundation grant (FDN388879 to J.-Y.M.). J.-Y.M. is a FRQS chair in genome stability.

## AUTHOR CONTRIBUTIONS

A.B.S., H.E. and E.V.D. conceived the study. A.B.S. performed experiments to characterize the role of XAB2 in seDSB repair; H.E. carried out the shRNA screen; L.P., M.-C.C, and K.N. performed experiments; P.V.N, B.K. and S.F. contributed to the screen and its analysis; P.V.N, A.B.S., H.E., M.-C.C., J.-Y.M. and L.P. carried out statistical data analysis; C.C.H.-M., P.C. and S.B. contributed cell lines for the study; S.P.N. provided material support and contributed to the screen analysis; J.-Y.M. and S.P.N. provided advice and supervision; P.C., J.-Y.M. and S.B. designed experiments and contributed to data analysis and interpretation. E.V.D. designed, performed and supervised the experiments and wrote the manuscript with comments from the authors.

## Competing interests

The authors declare no competing financial interests.

## Notes

### Competing Interest Statement

The authors have declared no competing interest.

