## Supplementary Figures and Legends for "XAB2 prevents abortive recombinational repair of replication-associated DNA double-strand breaks and its loss is synthetic lethal with RAD52 inhibition"

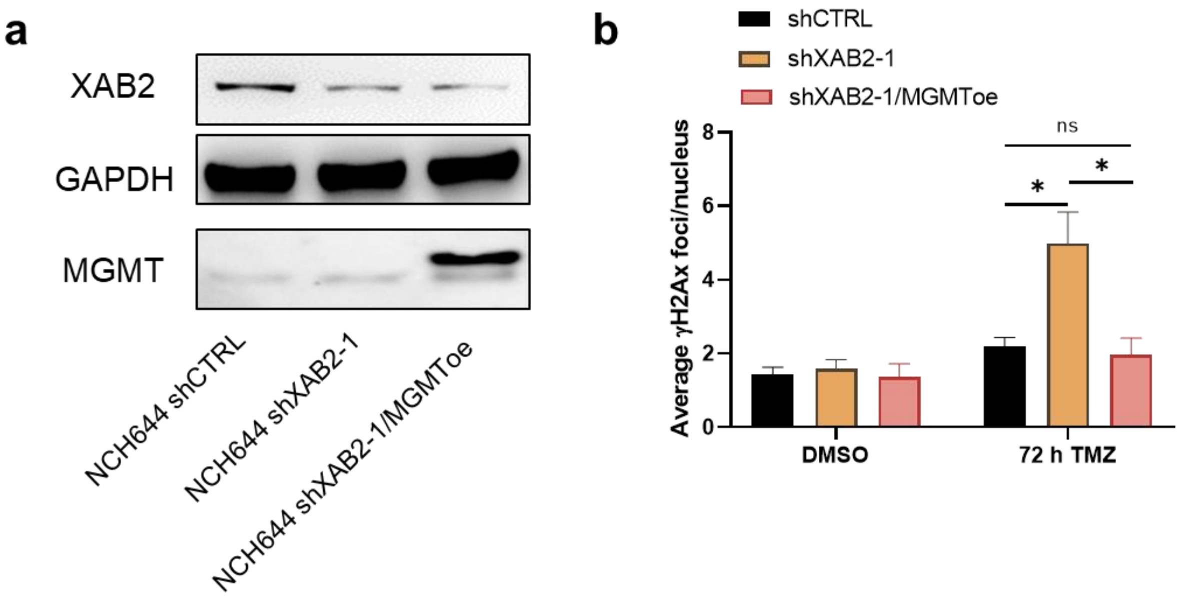


**Supplementary Fig. 1 | Ectopic MGMT expression prevents accumulation of TMZ associated seDSBs. a**, Immunoblot analysis of XAB2 and MGMT expression in NCH644 cells expressing shCTRL or shXAB2-1, as well as shXAB2-1-cells overexpressing MGMT (MGMToe). GAPDH was used as a loading control. **b**, Quantification of the average number of γH2Ax foci per nucleus in the indicated NCH644 cells. Error bar ± s.e.m. Differences between specified comparisons were assessed by a Mann-Whitney test and their significance is highlighted by stars (*P<0.05)(ns=non significant).

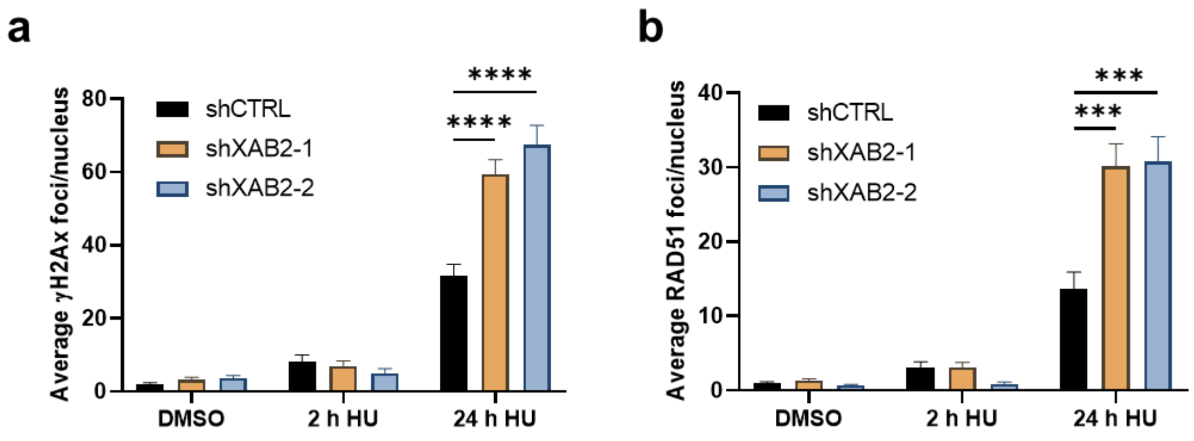


**Supplementary Fig. 2 | Loss of XAB2 leads to increased accumulation of seDSBs associated with collapsed replication forks. a-b**, U87 cells expressing the indicated shRNAs were incubated with hydroxyurea (HU) for 2 h or 24 h, followed by immunofluorescence analysis and quantification of γH2AX (**a**) and RAD51 (**b**) foci. Error bar ± s.e.m. Differences between specified comparisons were assessed by Kruskal-Wallis test and their significance is highlighted by stars (***P<0.001, ****P<0.0001).

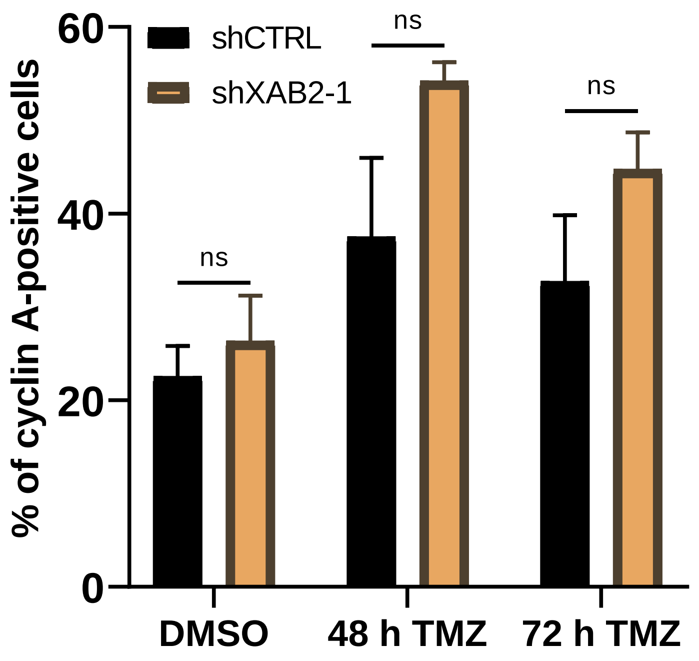

**Supplementary Fig. 3 | XAB2 depletion does not affects the cell cycle progression.** U87 cells expressing the indicated shRNAs were treated for 2 h with 15µM TMZ (or DMSO) and left to recover for 48 h or 72 h in drug-free medium before being processed for IF analysis of Cyclin A. The graph presents the percentage of Cyclin A-positive cells in the indicated conditions. Error bar ± s.e.m. Differences between specified comparisons were assessed by a Mann-Whitney test and their significance is highlighted by stars (*P<0.05; **P<0.01; ***P<0.001)(ns = non significant).

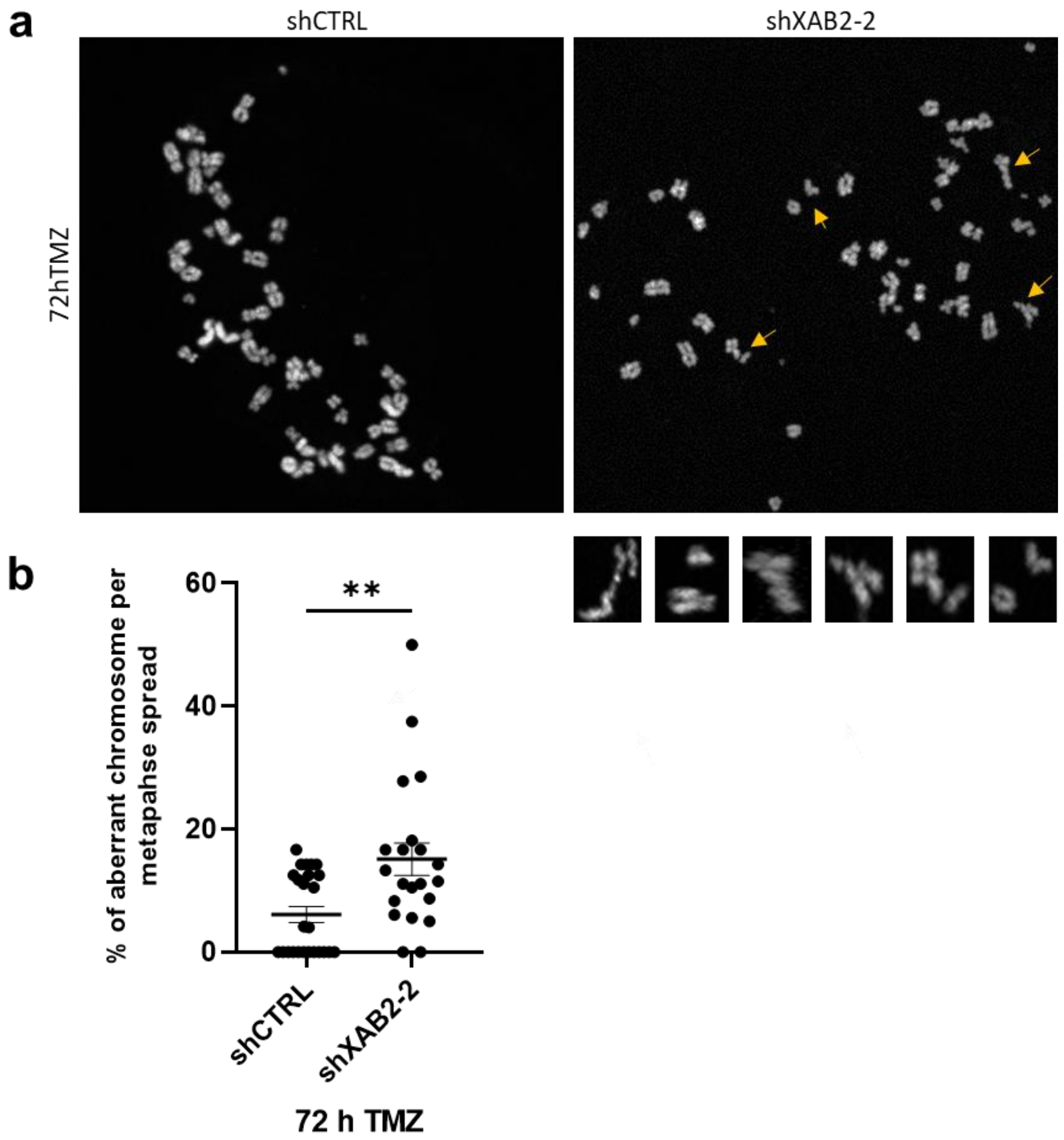


**Supplementary Fig. 4 | TMZ induces increased chromosomal aberrations in XAB2-depleted cells. a**-**b**, U87 cells with the indicated shRNAs were exposed to 15µM TMZ (or DMSO) for 2 h and left to recover in drug-free medium for 72 h, followed by metaphase spread preparation and microscopic analysis of chromosomal aberrations (**a**) and their quantification (**b**). Yellow arrows indicate cells for which enlarged images are presented in the lowest panel. Data are the average of n=2 biological replicates (30-50 cells/sample/experiment). Error bar ± s.e.m. Differences between samples were assessed by Mann-Whitney test and their significance is highlighted by stars (**$P<0.01$).

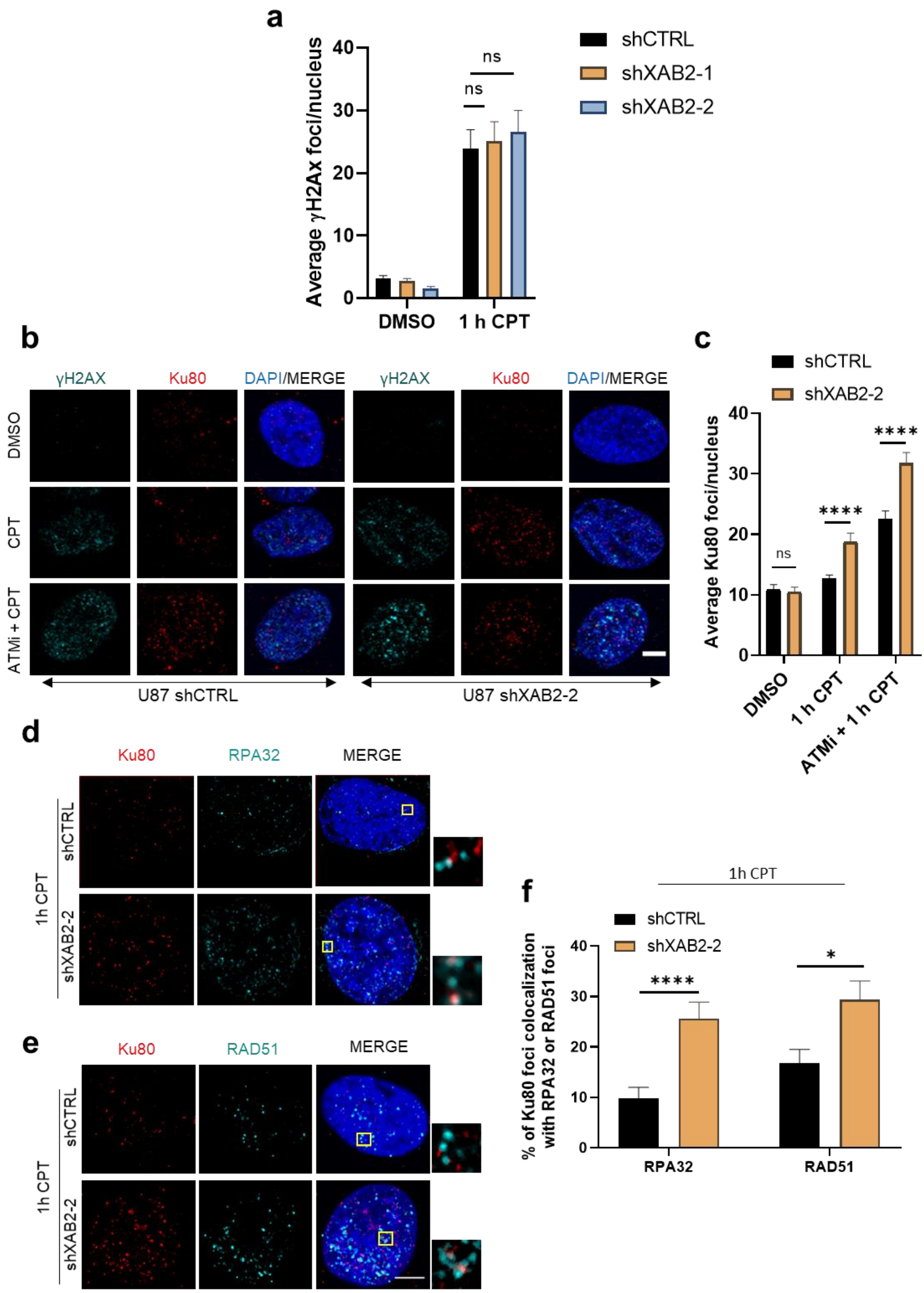
a
Average γH2Ax foci/nucleus
0
10
20
30
40
ns
ns
DMSO
1 h CPT
shCTRL
shXAB2-1
shXAB2-2
b
γH2AX
Ku80
DAPI/MERGE
γH2AX
Ku80
DAPI/MERGE
DMSO
CPT
ATMi + CPT
U87 shCTRL
U87 shXAB2-2
c
shCTRL
shXAB2-2
Average Ku80 foci/nucleus
0
10
20
30
40
ns
****
****
DMSO
1 h CPT
ATMi + 1 h CPT
d
Ku80
RPA32
MERGE
1h CPT
shCTRL
shXAB2-2
e
Ku80
RAD51
MERGE
1h CPT
shCTRL
shXAB2-2
f
1h CPT
shCTRL
shXAB2-2
% of Ku80 foci colocalization with RPA32 or RAD51 foci
0
10
20
30
40
****
*
RPA32
RAD51

**Supplementary Fig. 5 | XAB2 prevents Ku retention on seDSBs induced by campthotecin. a**, Graph representing the average number of γH2Ax foci per nucleus in U87 cells expressing the indicated shRNAs, following exposure to 1 µM CPT for 1 h and processing for IF. Error bar ± s.e.m. Differences between samples were assessed by Mann-Whitney test and their significance is highlighted by stars (ns = non-significant). **b**-**c**, Representative images of Ku80 foci (red) detected by immunofluorescence microscopy in control and XAB2-depleted cells exposed to 1 µM CPT for 1 h in the absence or presence of the ATMi KU-55933 (**b**) and related quantification (**c**). Cells were pre-treated with the ATMi or DMSO for 1 h prior to addition of CPT. Cells were also stained for γH2AX (cyan) and DNA was counterstained with DAPI (blue). Error bar ± s.e.m. Differences between samples treated with different drugs were assessed by Mann-Whitney test and their significance is highlighted by stars (****P<0.0001) (ns = non-significant). **d**, Representative immunofluorescence images used for the quantification of the number colocalized Ku80 (red) and RPA32 (cyan) foci in individual cells, following exposure of control and XAB2-depleted to 1 µM CPT for 1 h. The rightmost images present close up sections (rectangles) illustrating the extensive colocalization of RPA and Ku80 foci in XAB2-depleted cells compared to control cells. **e**, Same as in **d** for the analysis of Ku80 (red) and RAD51 (cyan) foci colocalization. **f**, Quantification of the frequencies of colocalized Ku80-RPA32 foci and Ku80-RAD51 foci in control and XAB2-depleted cells exposed to CPT, as assessed based on **d** and **e**. Scale bar= 5µm. Data are the average of n=3 biological replicates (30-50 cells/sample/experiment). Error bar ± s.e.m. Differences between samples were assessed by Mann-Whitney test and their significance is highlighted by stars (****P<0.0001) (ns = non-significant).

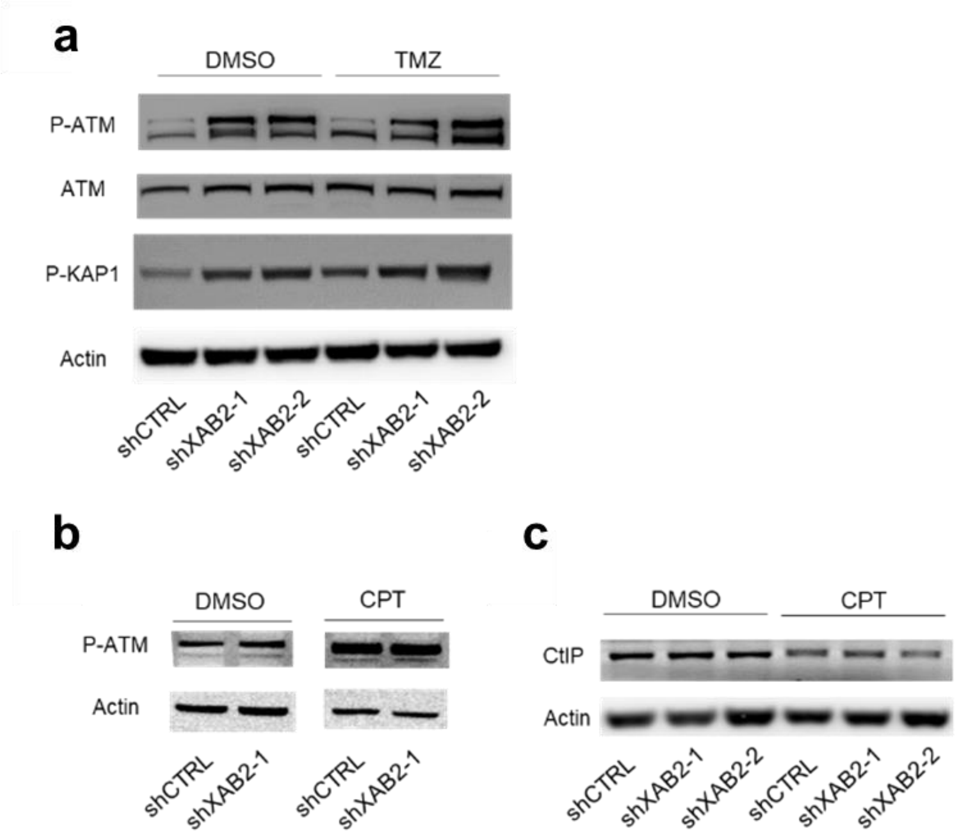


**Supplementary Fig. 6 | Impact of XAB2-depletion on ATM and CtIP phosphorylation induced by TMZ and CPT. a**, Extracts from control and XAB2-depleted U87 cells treated with 15 □M TMZ or (DMSO) for 2 h and allowed to recover for 48 h were subjected to immunoblotting analysis to examine ATM phosphorylation (P-ATM, using antibodies against pS1981) and KAP1 phosphorylation (P-KAP1, using antibodies against pS824). **b**, Immunoblotting analysis of P-ATM in extracts from control and XAB2-depleted cells following treatment with CPT (1 μM, 1h). **c**, Immunoblotting analysis of CtIP in extracts from control and XAB2-depleted cells following treatment with CPT. Note that exposure to CPT results in the hyper-phosphorylation of part of CtIP leading to the appearance of a doublet that does not resolve clearly. Such hyper-phosphorylation is not affected by XAB2 depletion.

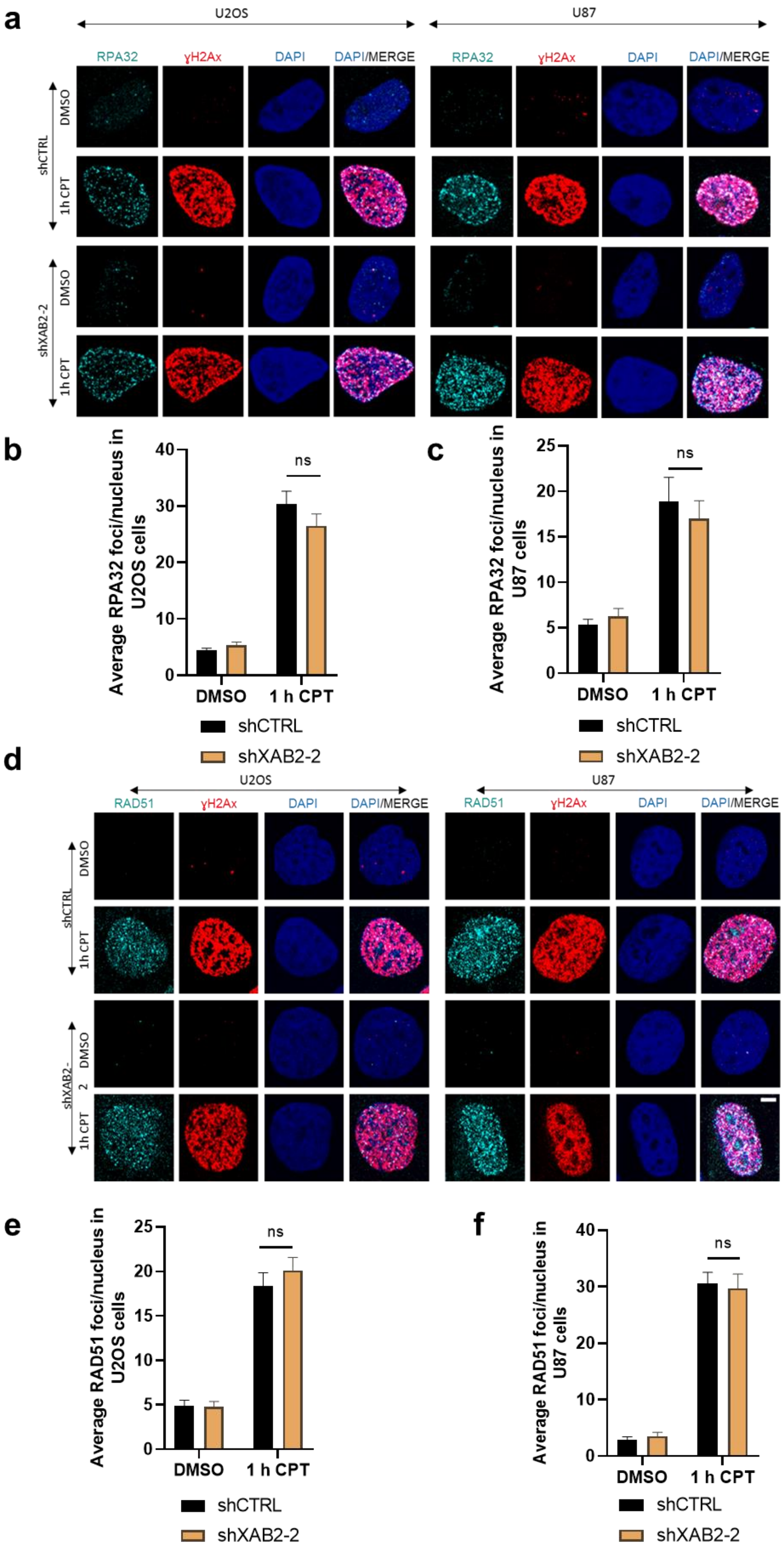
a
U2OS
U87
RPA32
γH2Ax
DAPI
DAPI/MERGE
shCTRL
DMSO
1h CPT
shXAB2-2
b
Average RPA32 foci/nucleus in U2OS cells
0
10
20
30
40
ns
DMSO
1 h CPT
shCTRL
shXAB2-2
c
Average RPA32 foci/nucleus in U87 cells
0
5
10
15
20
25
ns
DMSO
1 h CPT
shCTRL
shXAB2-2
d
U2OS
U87
RAD51
γH2Ax
DAPI
DAPI/MERGE
shCTRL
DMSO
1h CPT
shXAB2-2
e
Average RAD51 foci/nucleus in U2OS cells
0
5
10
15
20
25
ns
DMSO
1 h CPT
shCTRL
shXAB2-2
f
Average RAD51 foci/nucleus in U87 cells
0
10
20
30
40
ns
DMSO
1 h CPT
shCTRL
shXAB2-2

**Supplementary Fig. 7 | XAB2 depletion does not affect RPA32 and RAD51 foci formation in U2OS and U87 cells treated with CPT. a-c**, Representative immunofluorescence images of RPA32 (cyan) and gH2AX (red) foci in U2OS and U87 cells expressing shCTRL or shXAB2-2, following exposure to 1 μM CPT (or DMSO) for 1 h (**a**) and related quantification of RPA32 foci in U2OS (**b**) and U87 (**c**). **d-f**, Same as (**a-c**) for the analysis of RAD51 foci (cyan). Scale bar=5μm. Data are the average of n>2 biological replicates (30-50 cells/sample/experiment). Error bar ± s.e.m. Differences between samples treated with different drugs were assessed by Mann-Whitney test and their significance is highlighted by stars or ns (non-significant).

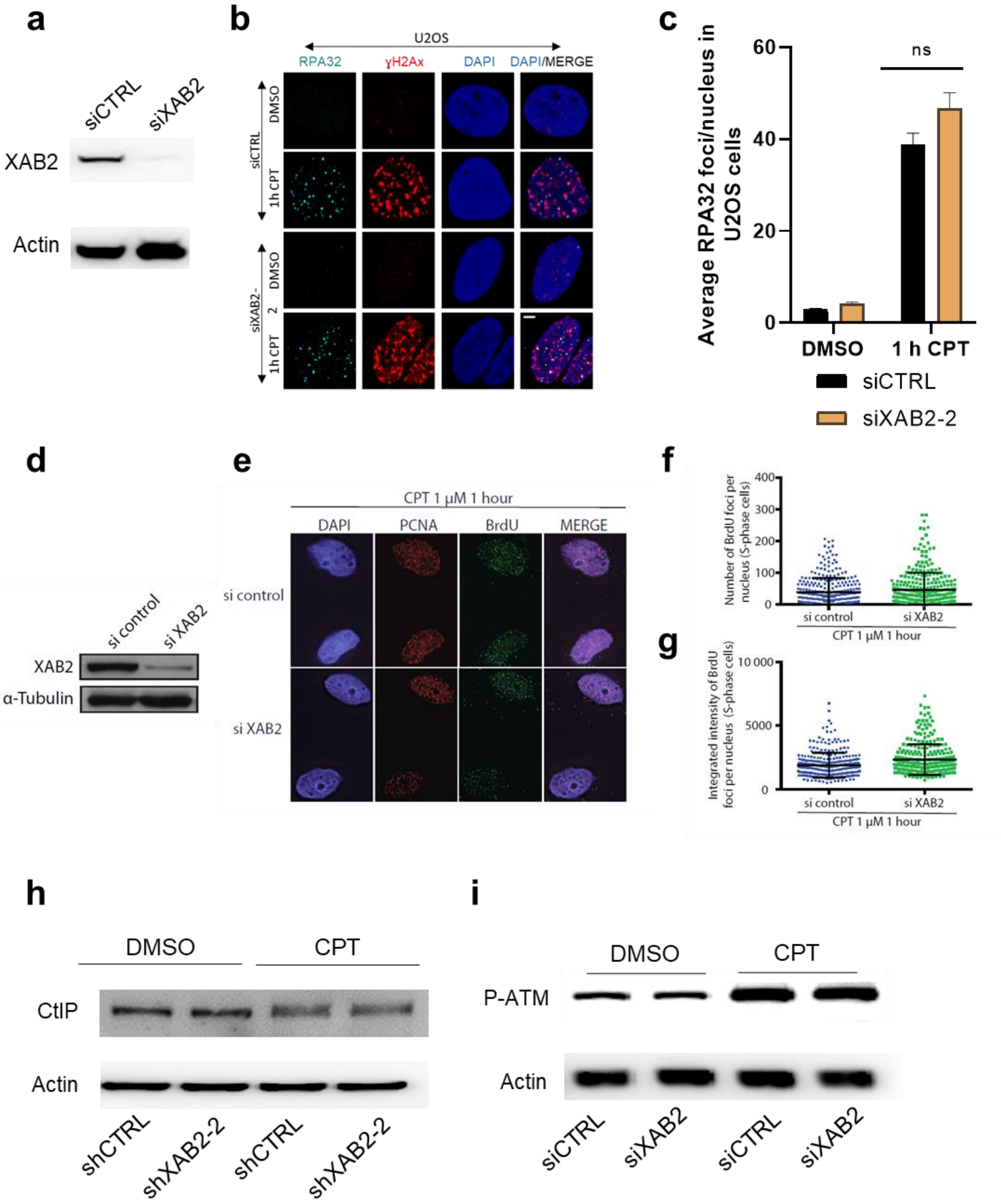
a
siCTRL
siXAB2
XAB2
Actin
b
U2OS
RPA32
γH2Ax
DAPI
DAPI/MERGE
siCTRL
DMSO
1h CPT
siXAB2-2
DMSO
1h CPT
c
Average RPA32 foci/nucleus in U2OS cells
ns
60
40
20
0
DMSO
1 h CPT
siCTRL
siXAB2-2
d
si control
si XAB2
XAB2
α-Tubulin
e
CPT 1 μM 1 hour
DAPI
PCNA
BrdU
MERGE
si control
si XAB2
f
Number of BrdU foci per nucleus (S-phase cells)
400
300
200
100
0
si control
si XAB2
CPT 1 μM 1 hour
g
Integrated intensity of BrdU foci per nucleus (S-phase cells)
10 000
5000
0
si control
si XAB2
CPT 1 μM 1 hour
h
DMSO
CPT
CtIP
Actin
shCTRL
shXAB2-2
shCTRL
shXAB2-2
i
DMSO
CPT
P-ATM
Actin
siCTRL
siXAB2
siCTRL
siXAB2

**Supplementary Fig. 8 | siRNA-mediated XAB2 depletion does not affect DNA end resection, and CtIP and ATM phosphorylation in U2OS treated with CPT.**

**a**-**c**, Analysis of RPA foci. **a,** Immunoblotting analysis of XAB2 depletion in U2OS cells treated with non-silencing siRNAs (siCTRL) or siRNAs targeting XAB2 (siXAB2). Actin was used as loading control. **b**-**c**, Representative immunofluorescence images of RPA32 (cyan) and gH2AX (red) foci in U2OS transfected with siCTRL or siXAB2 following exposure to 1 μM CPT (or DMSO) for 1 h (**b**), and related quantification of RPA32 foci (**c**). Scale bar= 5μm. Data are the average of n=3 biological replicates (30-50 cells/sample/experiment). Error bar ± s.e.m. Differences between samples treated with different drugs were assessed by Mann-Whitney test and their significance is highlighted by stars (ns = non-significant). **d**-**f**, Native BrdU assay to detect ssDNA. **d**, Western blotting to validate the knockdown of XAB2 in U2OS cells used in the assay. **e**, U2OS cells transfected with a control or XAB2 siRNA were incubated with BrdU for 24 h followed by treatment with 1 μM CPT for 1 h and a 1 h release period. The cells were then subjected to an immunofluorescence analysis against bromodeoxyuridine (BrdU) and PCNA. **f**, Quantification of the number of foci. The data show the mean ± s.d. The difference between the two conditions is not significant (Mann-Whitney U-test). **g**, Quantification of the integrated intensity of BrdU foci per nucleus. The data show the mean ± s.d. **** $p \leq 0.0001$ (Mann-Whitney U-test). **h**-**i**, Analysis of CtIP and ATM phosphorylation. **h**, Extracts from control and XAB2-depleted U20S cells treated with CPT for 1 h were subjected to immunoblotting analysis to examine CtIP phosphorylation (P-ATM, using antibodies against pS1981). Note that exposure to CPT results in the hyper-phosphorylation of part of CtIP leading to the appearance of a doublet that does not resolve clearly.  **i**,

Immunoblotting analysis of P-ATM in extracts from U2OS cells transfected with siCTRL or siXAB2, following treatment with CPT.
